# Task-general brain state dynamics relate to trial-to-trial performance and individual response strategy

**DOI:** 10.64898/2026.10.21.760632

**Authors:** Ayumu Yamashita, Masataka Sawayama, Kaoru Amano

## Abstract

Behavior fluctuates from moment to moment, yet individuals also show characteristic response strategies across tasks. Whether these two levels of variation relate to common brain dynamics is unknown. Using an inside-out approach across 10 cognitive tasks during fMRI, we identified recurring brain states and related their dynamics to behavior within and between individuals. Default mode network (DMN)–dominant and action mode network (AMN)–dominant states occurred in all tasks. Within individuals, momentary state identity (the state occupied at a given moment) tracked trial-to-trial performance. The AMN-dominant state was associated with slower, more variable responses across tasks, whereas its relationship to accuracy depended on the task. Across individuals, switching between the two states, but not time spent in either, showed cross-task consistency, and frequent switchers showed faster, less accurate responding. Thus, distinct aspects of the same task-general brain state dynamics, momentary state identity and state switching, relate to behavior at both levels.

## 1. Introduction

A central goal of systems and cognitive neuroscience is to understand how brain activity relates to behavior, in particular to performance in cognitive tasks. Traditionally, this has been studied with an outside-in approach: controlled stimuli or tasks are presented and the evoked neural responses are mapped onto specific cognitive functions to identify their neural mechanisms^1–3^. To understand cognitive functions at a general level that goes beyond any individual task, considerable effort has been devoted to organizing them into an explicit ontology of psychological constructs shared across tasks^4,5^. However, even within a single perceptual decision-making task, similar behavioral performance can arise from multiple brain activation patterns^6^. Across tasks, moreover, a construct defined in one paradigm is not guaranteed to engage the same psychological process in another, and comparison across studies therefore remains difficult.

A complementary, inside-out approach begins instead with the brain’s own ongoing activity and asks how that activity relates to behavior^7,8^. Ongoing activity can be summarized as transitions among a limited number of shared, whole-brain patterns, or brain states^8,9^. Prestimulus ongoing activity shapes perceptual decisions^10^, and brain states have been linked to cognition and behavior in task and naturalistic settings^11–13^. In our previous studies of a sustained attention task, transitions between two dominant brain states tracked differences in accuracy and reaction-time variability^14,15^. Because brain states are defined from the data rather than from predefined mental categories, the same states can be identified and compared across tasks even when psychological constructs are hard to equate^16^. Importantly, defining neural states independently of task-specific behavioral constructs makes it possible to ask whether the same brain state carries a common behavioral signature across tasks or has different behavioral consequences depending on context.

Recent studies have begun to identify shared brain states across diverse cognitive contexts^11,12,17^. Yet their behavioral significance has been examined at two largely separate levels. Studies of cross-task states have primarily emphasized between-individual differences, relating state occupancy or transition dynamics to fluid intelligence or cognitive control ability^12,17^. In contrast, within-individual studies have linked momentary state occupancy to trial-to-trial performance in sustained-attention tasks or to self-reported engagement during movie viewing^11,14,15^, but have generally been confined to a single task or setting. These levels of analysis are not interchangeable. A state metric that covaries with differences between people need not track performance fluctuations within a person, and a within-task state–behavior association need not generalize across tasks. Thus, it remains unknown which aspects of a shared state’s moment-to-moment behavioral signature are preserved across tasks, and which aspects of its dynamics are stable within individuals across tasks and relate to cross-task behavioral characteristics.

Here, we extended the inside-out approach across 10 cognitive tasks to bridge within- and between-individual levels of analysis. We first identified brain states separately in each task and integrated the resulting patterns into clusters that occur across tasks. We then examined within-individual coupling between state occupancy and trial-to-trial performance, and between-individual associations between state dynamics and cross-task behavioral characteristics. We further characterized event-related changes in state occupancy around demands for cognitive control. Finally, we examined state engagement during value-based decision-making and planning, asking whether occupancy tracked trial phase, task demand, and upcoming choice. These analyses asked which aspects of a shared state’s moment-to-moment behavioral signature generalize across tasks, which depend on task context, and whether occupancy or transition dynamics constitute stable individual characteristics related to behavior across tasks. Figure 1 provides an overview of the study design and main analyses.

**Figure 1:**
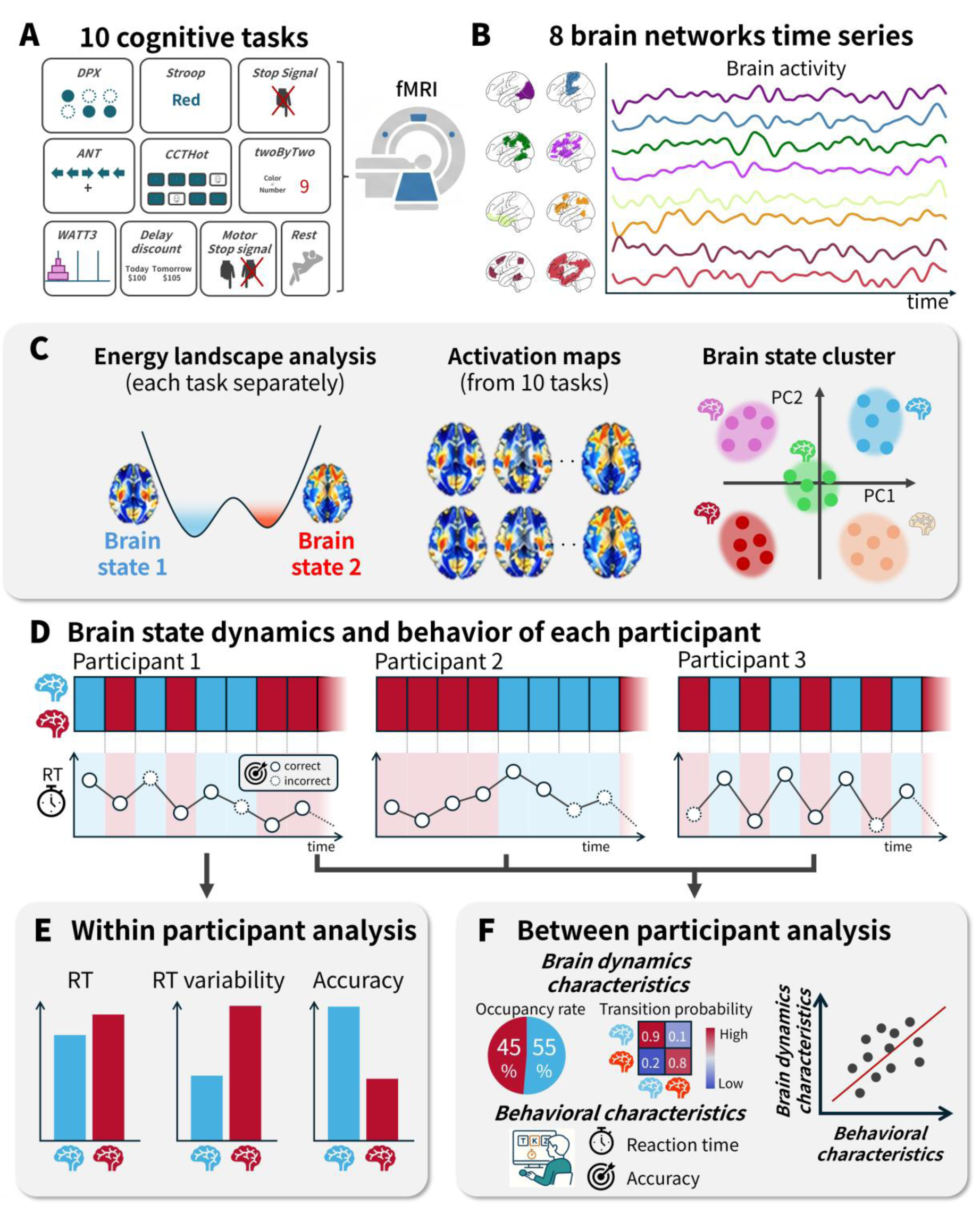
Study overview. (A) fMRI was acquired during 10 cognitive tasks (including rest). (B) Activity was summarized as eight network time series. (C) Brain states were identified with energy-landscape analysis in each task and integrated across tasks into brain state clusters by clustering voxel activation maps in the PC1–PC2 plane. (D) For each participant, the sequence of brain states and trial-to-trial behavior were obtained in each task. (E) Within participants, trial-to-trial performance was compared between brain state clusters (estimated across all participants with mixed-effects models). (F) Between participants, brain dynamics characteristics and behavioral characteristics were computed for each individual and related by across-participant correlation.

## 2. Results

### 2.1 Shared brain state clusters and temporal dynamics across tasks

To identify brain states that recur across cognitive contexts without imposing a common state structure a priori, we first estimated brain states independently within each of the 10 tasks. Each task yielded multiple brain states, defined as local minima of an energy landscape in which lower energy corresponds to more frequently occurring patterns of network activity (Methods; Figure 1 C), resulting in 36 task-specific brain states in total (Supplementary Fig. S1). We then represented each state as a voxelwise activation map and clustered these maps across tasks according to their spatial similarity, so that states discovered independently in different tasks could be grouped into common brain state clusters. Clustering these maps in the plane of the first two principal components (PC1– PC2; principal component analysis, PCA) produced seven brain state clusters (Clusters 0–6; Figure 2 A). The central result was that two of these clusters, Cluster 1 and Cluster 2, occurred in every cognitive task other than rest (Figure 2 E), even though brain states had been identified independently in each task. Cluster 1 showed default mode network (DMN)–dominant activity, whereas Cluster 2 showed action mode network (AMN)– dominant activity (Figure 2 D; Supplementary Figs. S2 and S3). Among subcortical regions, activity was strongest in the hippocampus in Cluster 1 and in the putamen in Cluster 2. Here, we use AMN to refer to a set of regions often assigned to the dorsal attention and ventral attention networks. Although other networks were also engaged in these states, we adopted this naming because recent work has emphasized a contrasting relationship between DMN and AMN activity^18^. The remaining clusters occurred only during rest or in subsets of tasks. A distinct resting-state pattern (Cluster 0) occurred exclusively during rest (Figure 2 E). Clusters 3 and 4 formed a second DMN–AMN opposed pair that occurred only in the value-based decision and planning tasks (discountFix, CCTHot, and WATT3), but differed from Clusters 1 and 2 in three respects (Figure 2 D; Supplementary Fig. S2). Frontoparietal control network A and B (FPCN-A and FPCN-B) were not opposed to each other, ventral attention and somatomotor activity were reversed relative to Clusters 1 and 2, and putamen involvement differed: putamen activity was negative in Cluster 1 but positive in Cluster 3, whereas Clusters 2 and 4 showed the opposite pattern (positive in Cluster 2 and negative in Cluster 4). Clusters 5 and 6 formed a third opposed pair, distinguished mainly by the presence versus absence of visual-network activity (Figure 2 D; Supplementary Fig. S2).

**Figure 2:**
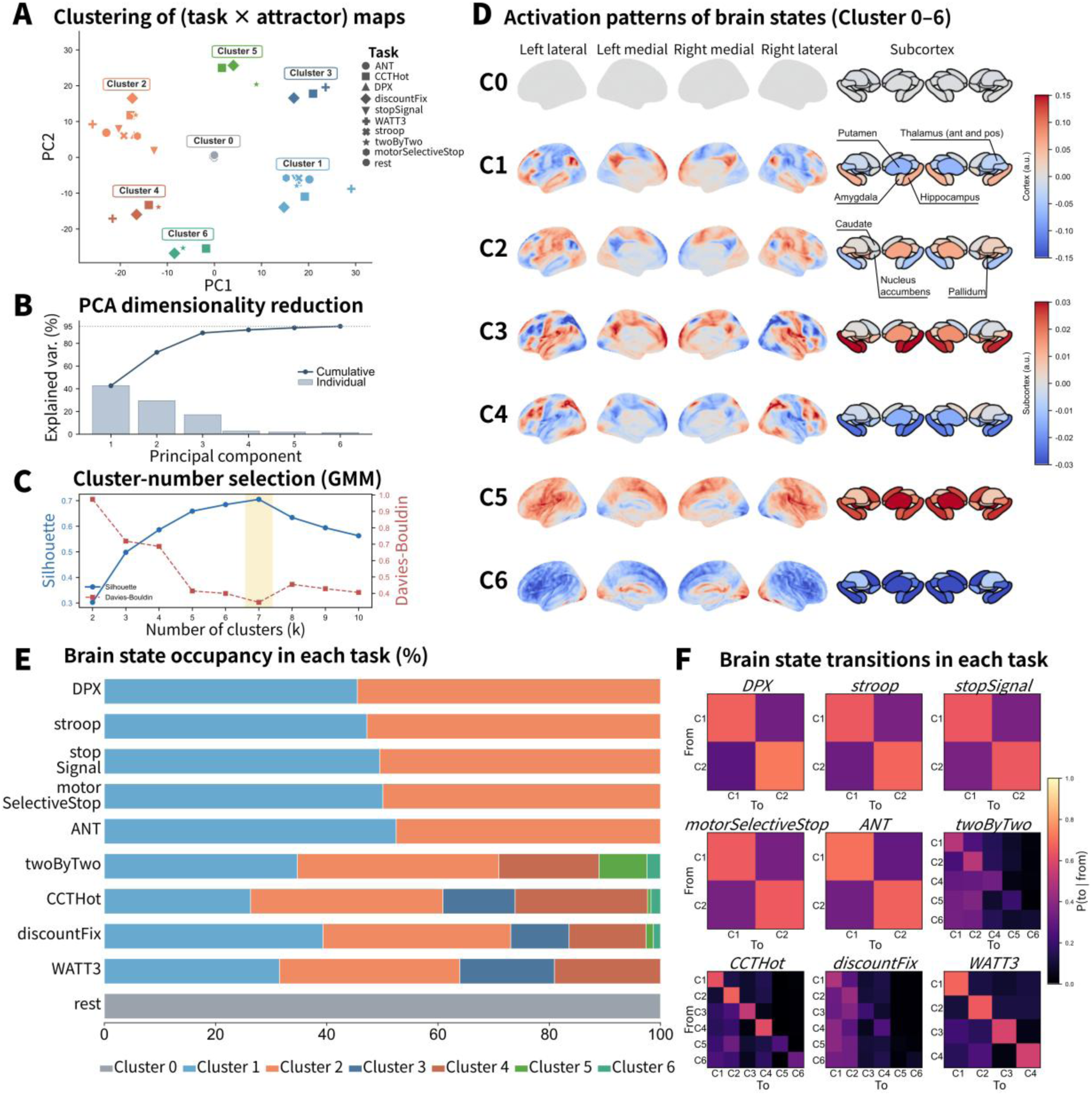
Cross-task identification and temporal dynamics of brain state clusters. (A) The 36 voxelwise activation maps obtained from task-specific brain states in 10 tasks were projected onto the first two principal components (PCs) and clustered using a Gaussian mixture model (GMM). Colors indicate brain state clusters and symbols indicate tasks. (B) Variance explained by each PC (bars) and cumulative explained variance (line); PC1 and PC2 explained 42% and 29% of the variance, respectively. (C) Silhouette and Davies– Bouldin scores for solutions with *k* = 2 − 10 clusters; *k* = 7, which maximized the silhouette score, is highlighted. (D) Mean cortical and subcortical activation maps for Clusters 0–6. Warm and cool colors indicate positive and negative activation, respectively; separate color scales are used for the cortex and subcortex. (E) Mean fractional occupancy of each brain state cluster in each task, averaged across participants and expressed as the percentage of time points. Each stacked bar sums to 100%. (F) Mean transition probabilities between brain state clusters at successive fMRI time points, averaged across participants. Rows indicate the cluster at the current time point (From), columns indicate the cluster at the next time point (To), and colors represent the conditional probability *P*(*To* ∣ *From*). Only clusters observed in each task are displayed. State-dynamics summaries included 90–99 participants per cognitive task and 106 participants for rest.

We next computed state occupancy and transition probabilities between all brain state clusters (Figure 2 E and F). Tasks in which only Clusters 1 and 2 were observed were mainly stimulus–response paradigms that required participants to respond as quickly and accurately as possible with a button press (ANT, Stroop, DPX, stopSignal, and motorSelectiveStop; see Methods for task descriptions). Clusters 5 and 6 had near-zero mean occupancy in most tasks (at most 1.7% outside twoByTwo, where Cluster 5 occupancy was 8.6%), and we did not include them in the subsequent behavioral analyses. Because the subsequent within- and between-individual analyses used reaction time (RT) and accuracy as behavioral measures, we restricted those analyses to Clusters 1 and 2, the clusters observed in the stimulus–response paradigms. Clusters 3 and 4 were examined in the value-based decision and planning analyses below.

The clustering solution itself was robust. PC1 and PC2 explained 42% and 29% of the variance across maps, respectively (71% cumulatively; Figure 2 B), and *k* = 7 was selected by the silhouette score (Figure 2 C). Hierarchical clustering of the same PC1–PC2 scores recovered the same seven clusters. Clustering in the 7-dimensional PCA subspace that retained 95% of the variance yielded a closely related partition (adjusted Rand index = 0.946). The silhouette-optimal solution had eight clusters only because one stopSignal map formed a cluster of its own, leaving the grouping essentially unchanged (Supplementary Methods, Clustering results and robustness).

### 2.2 Common brain state clusters show task-general associations with response speed but task-dependent associations with accuracy

We first asked, within individuals, how the brain state at the time of a trial (momentary state identity) related to behavior. Because Clusters 1 and 2 were the dominant states in the cognitive tasks characterized by repeated stimulus–response events, we compared Cluster 1 (DMN-dominant) with Cluster 2 (AMN-dominant). We found that trials associated with the AMN-dominant state (Cluster 2) were characterized by slower (linear mixed-effects model; state *β* = 0.024, SE = 0.003, *p* < 0.001) and more variable (state *β* = 0.023, SE = 0.003, *p* < 0.001) responses than trials associated with the DMN-dominant state (Cluster 1). The state × condition interaction was not significant for either measure (log-transformed RT [logRT], *χ*^2^(11) = 5.46, *p* = 0.91; RT variability [coefficient of variation, CV], *χ*^2^(11) = 7.92, *p* = 0.72; Figure 3 A, left and middle; Table 1). Across task conditions, the Cluster 2 − Cluster 1 contrast was consistently in the direction of slower and more variable responding (Figure 3 B, left and middle). This result indicates that the effect of brain state cluster on response speed and variability is task-general.

**Figure 3:**
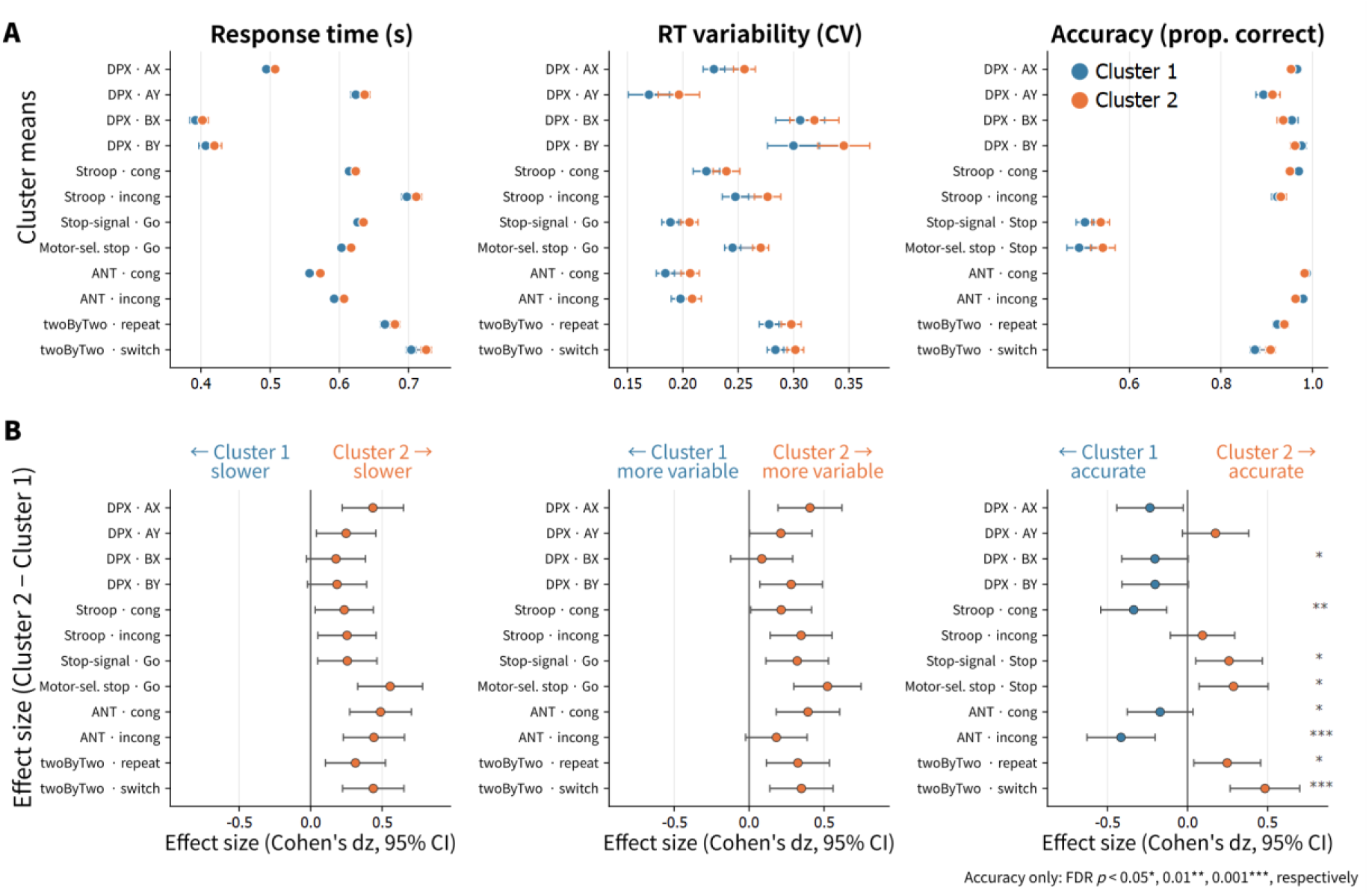
Within-individual associations between the two common brain states and task performance. Cluster 1 and Cluster 2 denote the default mode network (DMN)– and action mode network (AMN)–dominant states, respectively. Trials were assigned to the state observed 6 s after event onset. (A) Across-participant mean reaction time (RT), RT variability (coefficient of variation, CV), and accuracy for each state across 12 task conditions ( *n* = 89 –98 paired participants). RT is shown in seconds after back-transformation from logRT. For the stop-signal tasks, RT and CV were calculated from Go trials and accuracy from Stop trials. (B) Within-participant Cluster 2 − Cluster 1 differences (Cohen’s *d_z_*); positive values indicate slower responses, greater RT variability, or higher accuracy in Cluster 2. Asterisks denote FDR-adjusted paired Wilcoxon tests for accuracy (∗ *p_FDR_* < .05, ∗∗ *p_FDR_* < .01, ∗∗∗ *p_FDR_* < .001). Error bars show 95% CIs.

**Table 1:** Effect of brain state cluster on performance (linear mixed-effects models for logRT and CV; binomial generalized linear mixed model for accuracy). *β* > 0 indicates the direction in which Cluster 2 is slower / more variable / (in logit) more accurate.

| Outcome | state $\beta$ | SE | state $p$ | interaction $\chi^2$ (df) | interaction $p$ |
| --- | --- | --- | --- | --- | --- |
| Reaction time<br>(logRT) | 0.0244 | 0.0029 | < 0.001 | 5.46 (11) | 0.907 |
| RT variability (CV) | 0.0228 | 0.0031 | < 0.001 | 7.92 (11) | 0.720 |
| Accuracy (logit) | -0.1148 | 0.0285 | < 0.001 | 56.86 (11) | < 0.001 |

In contrast, for accuracy the state × condition interaction was significant (binomial generalized linear mixed model; state *β* = −0.115, SE = 0.029, *p* < 0.001; interaction *χ*^2^(11) = 56.9, *p* < 0.001; Figure 3 A, right). Because the interaction was significant, we followed up with paired Wilcoxon tests for each task condition, controlling the false discovery rate (FDR) with the Benjamini–Hochberg procedure. We found that the magnitude and direction of the state effect on accuracy differed across task conditions (significant in 8 of 12 task conditions, FDR < 0.05; Cohen’s *d_z_* in Figure 3 B, right). Thus, the Cluster 2 − Cluster 1 contrast in speed was consistent across tasks, whereas the same contrast for accuracy was predominantly task-dependent. Qualitatively, accuracy tended to be higher in Cluster 2 than in Cluster 1 in task conditions that appear to require greater cognitive control, such as the AY condition of DPX, the incongruent condition of the Stroop task, and stop trials in the stop-signal task (Figure 3 B; see Supplementary Methods, Cognitive tasks, for task and condition details).

### 2.3 State switching captures cross-task individual differences in the speed–accuracy trade-off

The preceding analyses showed that momentary state identity tracked trial-to-trial performance within individuals. We asked whether the dynamics of the two common states also contain a component that is consistent within individuals across tasks, and if so, what that component means for behavior.

#### 2.3.1 State switching shows greater cross-task individual consistency than state occupancy

We first asked which aspect of state dynamics reflects a stable individual characteristic (Figure 4 A and Table 2). Transitions between Cluster 1 and Cluster 2 dominated cognitive-task dynamics, so we focused on the transition probabilities between these two clusters. Furthermore, the C1→C2 and C2→C1 probabilities were highly correlated across participants and tasks (Spearman *r* = 0.944, *p* < 0.001, *n* = 856 participant-task observations), and we therefore averaged them as a single switch rate (C1↔C2).

**Figure 4:**
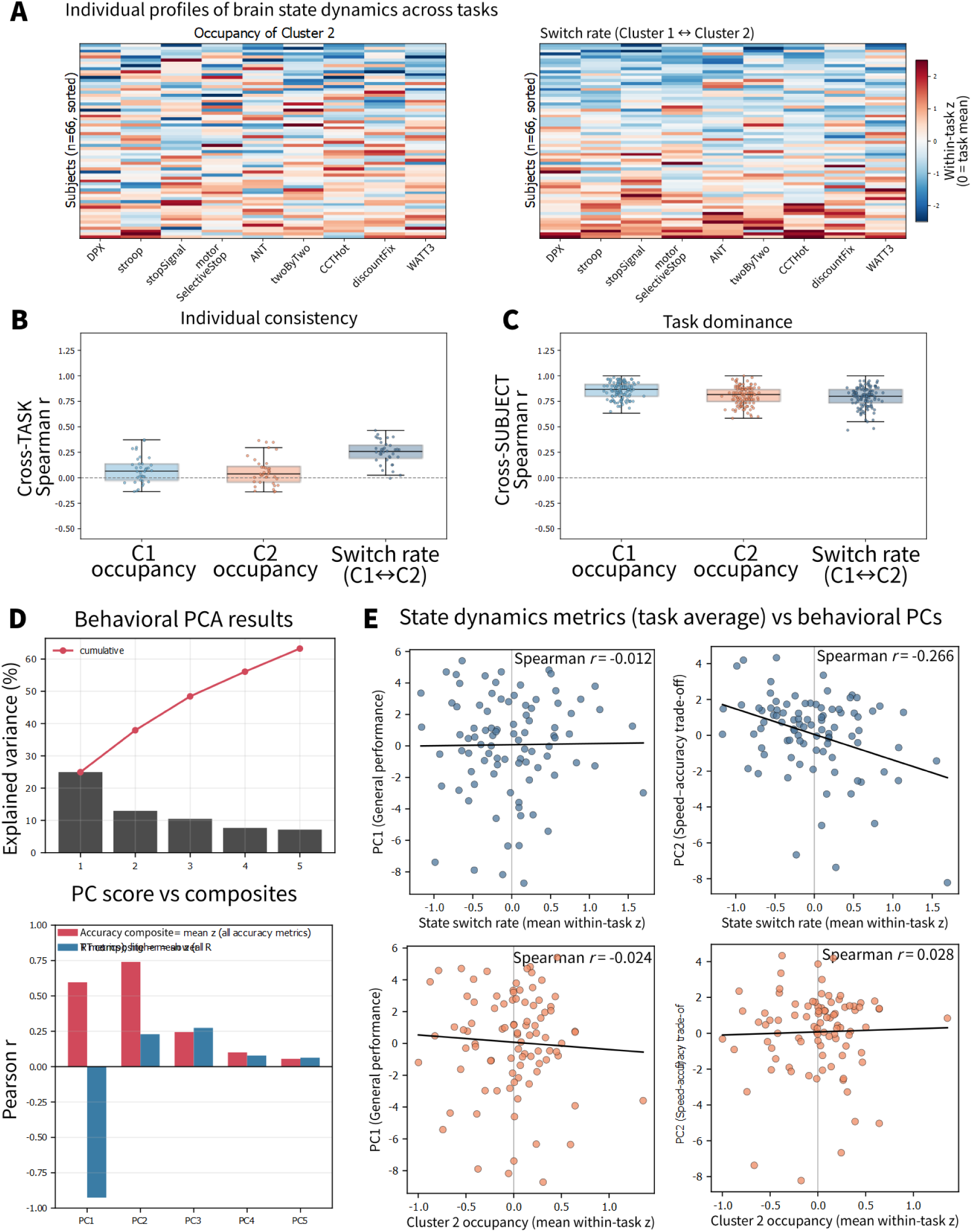
Cross-task consistency and behavioral correlates of individual state dynamics. Cluster 1 and Cluster 2 denote the DMN-and AMN-dominant states, respectively. (A) Individual profiles of within-task standardized Cluster 2 occupancy and bidirectional Cluster 1–Cluster 2 switch rate across nine non-rest tasks (*n* = 66 complete cases). (B) Spearman correlations across participants for all task pairs, indexing the cross-task consistency of individual differences. (C) Participant-profile correlations across the nine tasks, indexing task-related structure shared across participants. Boxes in B and C summarize the distribution of these pairwise correlations (median and interquartile range), with whiskers extending to 1.5 times the interquartile range. (D) Principal component analysis of accuracy and RT measures from six tasks (*n* = 94), showing explained variance and correlations with accuracy and RT composites. PC1 captured joint speed–accuracy performance, whereas PC2 captured the speed–accuracy trade-off. (E) Relationships of task-averaged switch rate and Cluster 2 occupancy with PC1 and PC2 (*n* = 91, at least four tasks); solid lines show ordinary least-squares fits for visualization.

**Table 2:** Participant-target and task-target ICC(3,1) of the state metrics. *n* = 66 participants, *k* = 9 non-rest tasks.

| State metric | participant-target |  | task-target |  |
| --- | --- | --- | --- | --- |
|  | ICC(3,1) | *p* | ICC(3,1) | *p* |
| State switch rate (C1-C2) | 0.299 | < 0.001 | 0.801 | < 0.001 |
| Cluster 1 occupancy | 0.068 | 0.0016 | 0.939 | < 0.001 |
| Cluster 2 occupancy | 0.015 | 0.222 | 0.937 | < 0.001 |

To distinguish systematic task-related differences from the cross-task consistency of individual differences, we estimated intraclass correlation coefficients (ICC(3,1)) in two complementary orientations. Task-target ICCs, which quantify whether tasks were ordered similarly for every participant, were high for all three state metrics: Cluster 1 occupancy (task-target ICC(3,1) = 0.939, *p* < 0.001), Cluster 2 occupancy (task-target ICC(3,1) = 0.937, *p* < 0.001), and the C1↔C2 switch rate (task-target ICC(3,1) = 0.801, *p* < 0.001; *n* = 66 participants, *k* = 9 non-rest tasks; Figure 4 C, Table 2). These high task-target ICCs indicate that both state occupancy and switching dynamics were strongly structured by task context and highly reproducible across participants.

Participant-target ICCs, which quantify whether participants were ordered similarly in every task, showed a different pattern (Figure 4 A). Cross-task individual consistency was minimal for occupancy. Cluster 2 occupancy did not show a significant participant-target ICC (ICC(3,1) = 0.015, *p* = 0.222), and Cluster 1 occupancy showed a statistically significant but very small ICC (ICC(3,1) = 0.068, *p* = 0.0016). By contrast, the switch rate showed a larger participant-target ICC (ICC(3,1) = 0.299, *p* < 0.001; Figure 4 B, Table 2). This result indicates that individuals retained some consistency in how frequently they switched between Clusters 1 and 2 across cognitive tasks.

Taken together, these complementary ICC analyses show that all three state metrics were strongly task-dependent, but that the switch rate additionally captured a cross-task individual component that was largely absent from occupancy.

#### 2.3.2 Switch rate is associated with a speed–accuracy trade-off

We next asked whether cross-task-consistent state dynamics relate to individual behavioral characteristics. Because switch rate showed greater cross-task individual consistency than occupancy (see above), we examined whether the task-averaged switch rate between Cluster 1 and Cluster 2 related to individual behavioral characteristics. As a control, we also examined occupancy. First, we summarized behavioral performance across tasks with principal component analysis (Figure 4 D). Higher scores on PC1 were associated with both higher accuracy and faster responding. We therefore interpreted PC1 as general performance. Higher scores on PC2 were associated with higher accuracy but slower responding, a speed–accuracy trade-off. This decomposition is consistent with prior reports that task-battery performance separates into ability and strategy dimensions^5,19,20^. We then asked whether the switch rate relates to general performance (PC1) or a speed–accuracy trade-off (PC2) (Figure 4 E, Supplementary Table S5). We found that the task-averaged switch rate was negatively correlated with behavioral PC2 (the speed–accuracy trade-off; Spearman *r* = −0.266, *p* = 0.011, FDR-adjusted *p* = 0.022, *n* = 91; Figure 4 E top right; Supplementary Table S5), but not with PC1 (general performance; *r* = −0.012, *p* = 0.91, FDR-adjusted *p* = 0.91; Figure 4 E top left). We also found no correlation between Cluster 1 or Cluster 2 occupancy and either PC1 or PC2 (Cluster 1: PC1, *r* = −0.009, *p* = 0.93, FDR-adjusted *p* = 0.93; PC2, *r* = 0.090, *p* = 0.40, FDR-adjusted *p* = 0.79; Cluster 2: PC1, *r* = −0.024, *p* = 0.82 ; PC2, *r* = 0.028, *p* = 0.79 ; both FDR-adjusted *p* = 0.82 ; Supplementary Table S5; Cluster 2 in Figure 4 E bottom). As a robustness check, restricting the analysis to participants with all nine non-rest tasks yielded the same association with PC2 (Spearman *r* = −0.282, *p* = 0.022, FDR-adjusted *p* = 0.043, *n* = 66). This result indicates that a higher switch rate was associated with behavior favoring speed over accuracy.

Taken together with the within-individual results, these findings indicate that distinct aspects of the same brain state dynamics related to behavior within and between individuals. Momentary state identity tracked trial-to-trial fluctuations in performance, whereas switching between the two states tracked stable individual differences in response strategy across tasks.

### 2.4 AMN-dominant state occupancy increases with cognitive-control demand and differs between successful and failed stopping

We next examined whether the occupancy of the two common states changed with moment-to-moment cognitive control demands and response-inhibition outcomes using the peri-event contrasts described in the Methods (Figure 5). Because Clusters 1 and 2 were the only states observed in these tasks and their occupancy time courses were therefore complementary, here we report only Cluster 2. Using cluster-based permutation tests of the within-participant condition difference, we found that Cluster 2 occupancy was higher under high-control than low-control conditions in the Stroop task (incongruent versus congruent; 3.4–6.8 s after the event; cluster statistic = 24.8, *p* < 0.001, *n* = 96; Figure 5 A left) and in the cued task-switching task (twoByTwo; switch versus stay; 2.72–4.76 s; cluster statistic = 11.1, *p* = 0.002, *n* = 92; Figure 5 A right). In the ANT, we did not find a significant high-control (incongruent) versus low-control (congruent) difference (peak at 4.08 s after the event; cluster statistic = 2.3, *p* = 0.213, *n* = 93; Figure 5 A, middle), but a similar tendency was observed. These findings are consistent with transient recruitment of the AMN-dominant state under increased cognitive-control demands.

**Figure 5:**
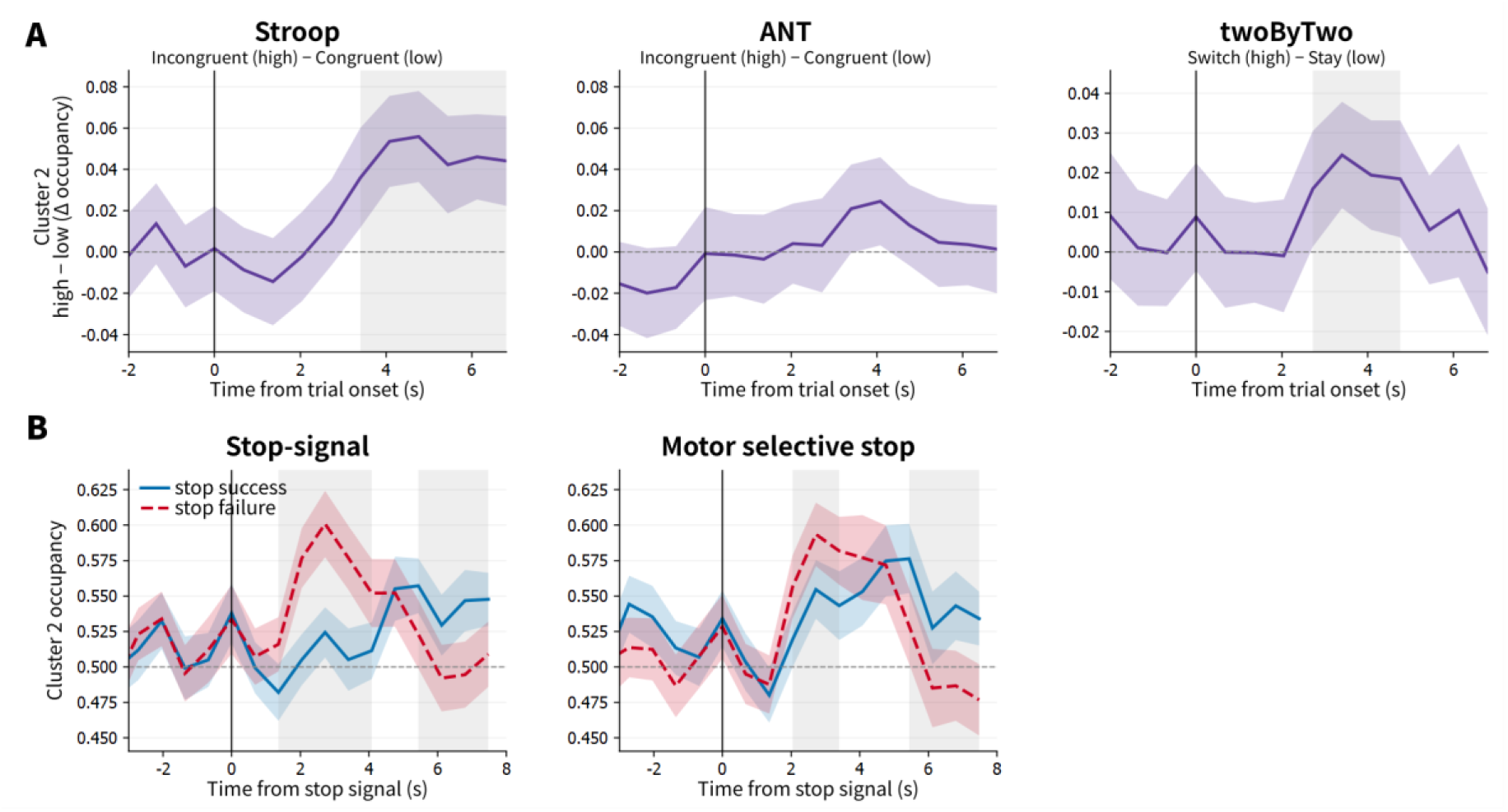
Event-related temporal dynamics of the AMN-dominant state (Cluster 2). (A) Trial-onset–aligned differences in Cluster 2 occupancy between high-and low-control trials: incongruent minus congruent trials in Stroop (*n* = 96) and ANT (*n* = 93), and switch minus stay trials in twoByTwo ( *n* = 92 ). (B) Stop-signal–aligned Cluster 2 occupancy for successful and failed stop trials in stopSignal ( *n* = 93 ) and motorSelectiveStop ( *n* = 87 ). Lines show across-participant means and colored bands show 95% CIs; solid blue and dashed red lines in B denote successful and failed trials, respectively. Vertical black lines mark event onset, horizontal dashed lines mark zero difference in A and occupancy = 0.5 in B, and gray shading marks significant temporal clusters from paired cluster-based permutation tests corrected across time. Time courses are shown without a fixed hemodynamic shift.

In the two stop-signal tasks, we found that Cluster 2 occupancy was higher on failed than successful stop trials from 1.36–4.08 s after the event in stopSignal (cluster statistic = 20.7, *p* < 0.001, *n* = 93; Figure 5 B left) and lower on failed than successful trials from 5.44–7.48 s (cluster statistic = −10.6, *p* = 0.013, *n* = 93; Figure 5 B left). We found a similar early positive difference in motorSelectiveStop (2.04–3.4 s; cluster statistic = 7.1, *p* = 0.046, *n* = 87 ; Figure 5 B right) and a late negative difference (5.44–7.48 s; cluster statistic = −11.8, *p* = 0.0098, *n* = 87; Figure 5 B right). Together, these results indicate that successful and failed stopping were associated with distinct temporal profiles of AMN-dominant state occupancy that were consistent across the two stop-signal tasks.

### 2.5 Brain state clusters in value-based decision and planning paradigms

Finally, as a complementary analysis to the executive-performance links above, we examined how brain state clusters related to decision and planning variables in discountFix, CCTHot, and WATT3 (Figure 6).

**Figure 6:**
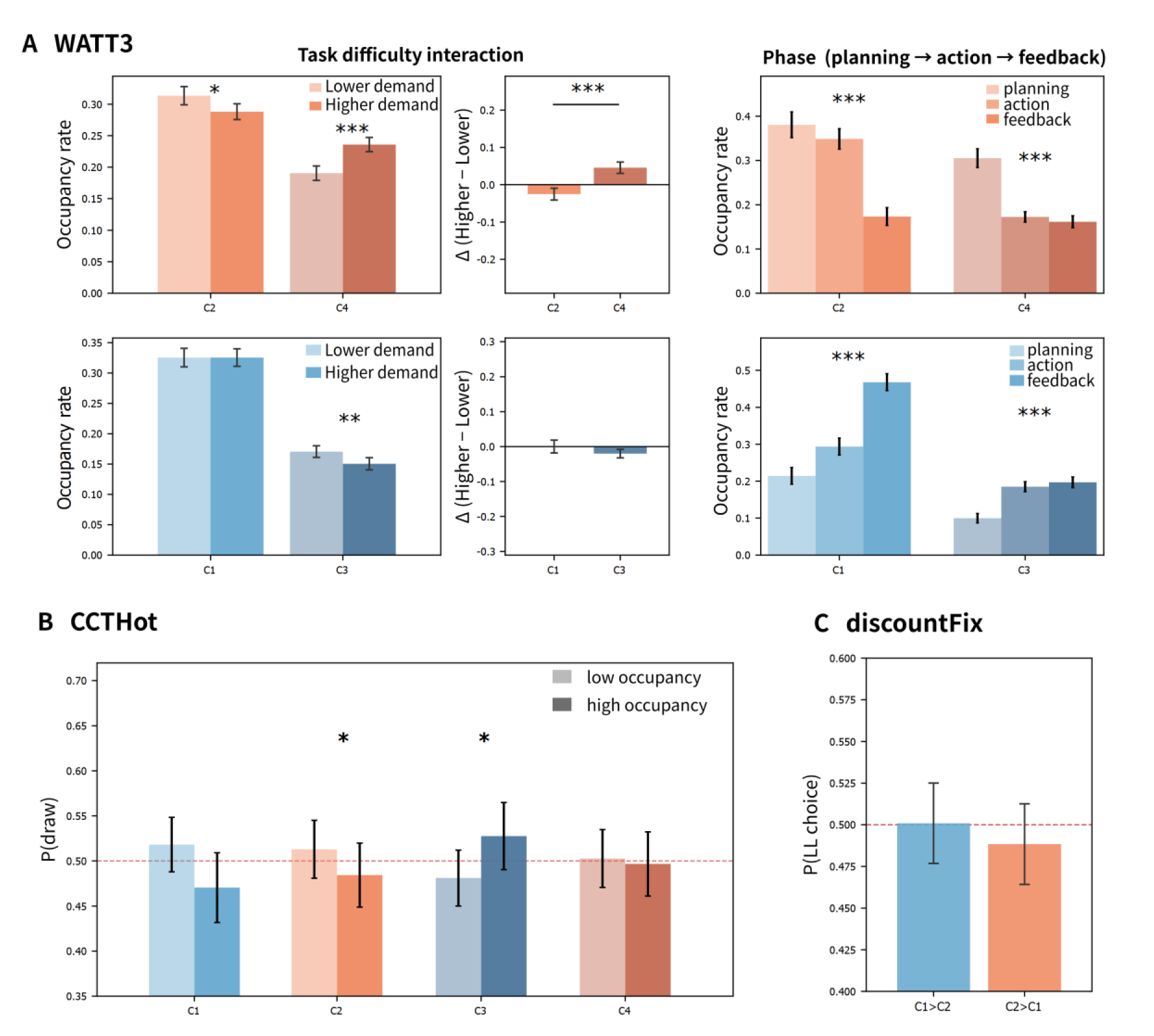
Brain state occupancy during planning and value-based decision making. Clusters 1 and 3 are DMN-dominant states, and Clusters 2 and 4 are AMN-dominant states. (A) WATT3 (*n* = 96): occupancy in lower-demand problems without versus higher-demand problems with an intermediate move (left), higher-minus-lower demand contrasts (center), and occupancy during planning, action, and feedback (right). Brackets compare clusters within each network pair. (B) CCTHot (*n* = 89): probability of drawing on expected-value-matched final decisions at lower versus higher cluster occupancy during the preceding seven TRs. (C) discountFix ( *n* = 94 ): probability of choosing the larger-later option according to whether Cluster 1 or Cluster 2 occupancy was greater during the preceding seven TRs. Bars in B and C are descriptive; inference used continuous occupancy in participant-random-intercept logistic mixed models. Unbracketed and bracketed asterisks in A denote within-cluster effects and between-cluster interactions, respectively; asterisks in B denote FDR-adjusted occupancy effects. Error bars show 95% CIs. Red dashed lines indicate a probability of 0.5. All analyses used a 6-s hemodynamic shift.

In WATT3, a towers planning task, we compared the occupancy of Clusters 1–4 between lower-demand problems (solvable without an intermediate move) and higher-demand problems (requiring an intermediate move). The effect of planning demand differed between clusters even within the same dominant-network family (Clusters 1 and 3, DMN-dominant; Clusters 2 and 4, AMN-dominant; Figure 6 A left). Among the AMN-dominant states, Cluster 4 occupancy was higher under higher demand (*β* = 0.045, SE = 0.007, *p* < 0.001), whereas Cluster 2 occupancy was lower (*β* = −0.026, SE = 0.007, *p* < 0.001). Among the DMN-dominant states, Cluster 3 occupancy was lower under higher demand (*β* = −0.021, SE = 0.007, *p* = 0.0021), whereas Cluster 1 was unchanged (*β* = 0.002, SE = 0.007, *p* = 0.72; *n* = 96). Demand-by-cluster interactions confirmed these within-pair differences (AMN-dominant pair, *β* = −0.071, SE = 0.010, *p* < 0.001 ; DMN-dominant pair, *β* = 0.024, SE = 0.010, *p* = 0.015). These results indicate that sensitivity to planning demand was cluster-specific and could not be explained solely by whether a state was DMN-or AMN-dominant. In particular, the selective increase in Cluster 4 suggests that this AMN-dominant state was more closely related to demanding planning than the common AMN-dominant state (Cluster 2). Cluster 4 differed from Cluster 2 mainly in FPCN-A, FPCN-B, and putamen activity (Figure 2 D), a difference we consider in the Discussion.

We also found phase-dependent occupancy (planning, action, and feedback phases) for all four clusters (Wald tests; Cluster 1, *χ*^2^(2) = 1005, *p* < 0.001; Cluster 2, *χ*^2^(2) = 753, *p* < 0.001 ; Cluster 3, *χ*^2^(2) = 164, *p* < 0.001 ; Cluster 4, *χ*^2^(2) = 359, *p* < 0.001 ; Figure 6 A right). Cluster 2 and Cluster 4 occupancy was highest during planning and lowest during feedback, whereas Cluster 1 and Cluster 3 occupancy was lowest during planning and highest during feedback. This temporal organization indicates that state occupancy tracked different stages within a trial, with the AMN-dominant states preferentially expressed during planning and action and the DMN-dominant states becoming more prominent after planning, particularly during feedback.

In CCTHot, a sequential risk-taking card task, we asked whether brain state occupancy immediately preceding a decision was related to whether participants stopped or continued. For this purpose, we compared the pre-decision occupancy of each cluster between round-final stop decisions and continue decisions matched within participants for expected value (846 pairs, *n* = 89; Figure 6 B). We found that higher pre-decision occupancy of Cluster 3 was associated with a higher probability of continuing (*β* = 0.547, SE = 0.199, *p* = 0.006, FDR *p* = 0.024, odds ratio [OR] = 1.73), whereas higher occupancy of Cluster 2 was associated with a lower probability of continuing (*β* = −0.214, SE = 0.090, *p* = 0.018, FDR *p* = 0.036, OR = 0.81). We did not find significant associations for Clusters 1 or 4 (Cluster 1, *β* = −0.179, SE = 0.155, *p* = 0.25; Cluster 4, *β* = 0.121, SE = 0.133, *p* = 0.36; 1,692 decisions). Because stop and continue decisions were matched on expected value, these results indicate that pre-decision occupancy of Clusters 2 and 3 was associated with the upcoming choice and could not be attributed simply to differences in expected value.

In discountFix, an intertemporal choice task between a smaller-sooner (SS) and a larger-later (LL) reward, we asked whether brain state occupancy immediately preceding a choice was related to which option participants chose. We focused on mid trials, in which LL and SS options were nearly equally preferred and choice was therefore expected to be most difficult. We restricted this analysis to Clusters 1 and 2 because Clusters 3 and 4 did not provide enough trials for reliable estimation. We then tested whether the probability of an LL choice depended on the balance between Cluster 1 and Cluster 2 occupancy preceding the response (Cluster 1 minus Cluster 2 occupancy over ≈ 4.8 s, seven fMRI volumes [TRs]; Figure 6 C). As a result, we did not find an association (*β* = −0.053, SE = 0.079, *p* = 0.50, OR = 0.95; 3,588 trials, *n* = 94).

Together, these analyses indicate that the behavioral significance of brain state occupancy depended on task context. In WATT3, occupancy tracked planning demand and task phase, with dissociations even between network-matched clusters. In CCTHot, pre-decision occupancy of Clusters 2 and 3 was associated with whether participants stopped or continued after decisions were matched on expected value, whereas in discountFix the Cluster 1–Cluster 2 balance was not associated with intertemporal choice. These findings suggest that brain state occupancy is not uniformly related to decision behavior but instead reflects the specific demands and temporal structure of each paradigm.

## 3. Discussion

In this study, we used an inside-out approach to examine how shared brain state dynamics relate to behavior within and between individuals across multiple tasks. Two states, a DMN-dominant state (Cluster 1) and an AMN-dominant state (Cluster 2), occurred across all non-rest tasks. Within individuals, Cluster 2 was associated with slower and more variable responses than Cluster 1 across task conditions, whereas the association with accuracy depended on task condition. Across individuals, both occupancy and switch rate were strongly structured by task context, and the switch rate showed a modest cross-task individual component. Individuals with a higher switch rate favored speed over accuracy. DMN- and AMN-dominant variants (Clusters 3 and 4), whose activity patterns differed from Clusters 1 and 2, also occurred in the planning and value-based decision paradigms. Relationships between state occupancy and task variables differed across these paradigms. Together, these findings indicate that distinct aspects of the same task-general brain state dynamics relate to behavior within and between individuals. Momentary state identity tracked trial-to-trial fluctuations in performance, whereas switching between states tracked stable individual differences in response strategy across tasks. Task context shaped which states occurred and what their occupancy meant for behavior.

### 3.1 Task-general and task-dependent behavioral associations of the two common states

We first asked whether the contrast between the task-general DMN-dominant and AMN-dominant states (Clusters 1 and 2) carried a common behavioral meaning across tasks and behavioral measures. Previous studies have shown that task-specific activity patterns can be organized into shared large-scale configurations that reappear across rest, tasks, and contexts^11,12,17,21,22^, and these earlier studies have often linked such states to between-individual ability^12^ or to performance within a single paradigm^11,14,23^. However, this prior work left open whether a shared state carries the same functional meaning across diverse tasks and across different behavioral readouts. Here, across the analyzed task conditions, associations of the DMN-dominant versus AMN-dominant contrast with response speed and variability were relatively consistent, whereas associations with accuracy were task-condition-dependent. The consistent associations with response speed and variability established a common functional reference across tasks. With this common reference established, the task-dependent accuracy effects demonstrate that the same states can support different behavioral outcomes across contexts. This dissociation is difficult to establish with an outside-in approach, in which brain activity is identified from behavioral task performance, and to our knowledge the present inside-out design is the first to reveal it.

### 3.2 Interpretation of the two common states as automatic and controlled modes

These state–behavior associations raise the question of what cognitive modes the two common states reflect. Prior work has attributed a range of functions to the AMN and the DMN^18,24^. Among these proposals, we interpret the AMN-dominant state (Cluster 2) as reflecting engagement of cognitive control and the DMN-dominant state (Cluster 1) as a relatively automatic mode of processing^25–27^. Four observations from our data support this interpretation. First, being in the AMN-dominant state was associated with slower and more variable responding. This pattern is consistent with an effortful-control interpretation^28,29^. Second, accuracy tended to be higher in the AMN-dominant state than in the DMN-dominant state precisely in conditions that appear to require greater cognitive control, such as the AY condition of DPX, the incongruent condition of the Stroop task, and stop trials in the stop-signal tasks (Figure 3 B). Third, AMN-dominant state occupancy was higher under high-control than low-control conditions in Stroop and twoByTwo (Figure 5). On this account, the consistent associations with response speed and variability reflect a general cost of engaging control, whereas the benefit of the AMN-dominant state for accuracy emerges selectively in conditions whose requirements call for control^30,31^. Fourth, hippocampus activity was strongest in the DMN-dominant state, consistent with a memory-based automatic-processing interpretation^27^, and putamen activity was strongest in the AMN-dominant state, consistent with an action-related control interpretation^18^ (Figure 2 D). Nevertheless, because control demand was not directly tested as a moderator and our data are correlational, these results do not establish automatic versus controlled processing as the functional meaning of the two states. This hypothesis could be tested in future work by parametrically manipulating control demand within a task and examining whether state occupancy tracks the manipulation.

### 3.3 Individual differences in state dynamics and response strategy

Having established the within-individual functional distinction between the task-general DMN-dominant and AMN-dominant states, we next asked whether their dynamics also explain cross-task behavioral differences between individuals. Both occupancy and switch rate were strongly structured by task context, but the cross-task individual component was more evident in switch rate than in occupancy. We summarized behavioral performance across tasks into two principal components, general performance (PC1) and the speed– accuracy trade-off (PC2). The task-averaged switch rate was negatively associated with the speed–accuracy trade-off component, showing that individuals who switched more frequently favored speed over accuracy. No association with general performance was detected, and task-averaged occupancy of the DMN-dominant and AMN-dominant states was not associated with either behavioral component. The separation of general performance from the speed–accuracy trade-off accords with prior work distinguishing ability and strategy dimensions^5,19,20^. Thus, switching, rather than the average time spent in either state, captured an individual difference in the speed–accuracy trade-off.

Under the interpretation developed above, switch rate reflects how frequently an individual alternates between automatic and controlled modes of processing. We propose that frequent alternation reduces sustained engagement of the controlled mode and thereby favors faster responding at the cost of accuracy. This account connects the within-individual functional meaning of the two states to a between-individual difference in response strategy.

Our findings extend recent evidence that occupancy of a shared multiple-demand, task-positive state predicts task-specific cognitive-control ability across paradigms and shows cross-task consistency^12^. That work examined the shared state at the between-individual level, relating task-specific occupancy to task-specific control measures within each paradigm. The present study differs in two respects. First, we related task-averaged state metrics to behavioral components derived across tasks, asking which aspect of state dynamics constitutes a task-general individual characteristic. In our heterogeneous task battery, occupancy was dominated by task context, whereas switch rate captured the behaviorally relevant cross-task individual component. These findings point to complementary roles for the two metrics: occupancy captures how strongly a task recruits a state and therefore relates to ability within a paradigm, whereas switching captures a cross-task disposition closer to response strategy. Second, we examined the same two task-general states at both the within- and between-individual levels in the same participants. Momentary state identity tracked trial-to-trial fluctuations in performance, and switching between the same states tracked stable individual differences in response strategy across tasks. Because both associations were obtained for the same states, the within-individual results, which supported an interpretation of the two states as automatic and controlled modes, gave the between-individual switch rate a concrete functional meaning as the frequency of alternation between these modes.

### 3.4 State variants specific to planning and value-based decision making

Task context shaped not only the dynamics of the two common states but also the repertoire of states expressed. The planning and value-based decision paradigms additionally expressed Clusters 3 and 4, a second pair of DMN- and AMN-dominant states (Figure 6). These additional states allowed us to determine whether states within the same dominant-network family nevertheless serve distinct functional roles. Relative to Clusters 1 and 2, Clusters 3 and 4 were distinguished mainly by a dissociation between FPCN-A and FPCN-B and by a reversed pattern of putamen involvement (Figure 2 D).

In WATT3, occupancy varied with task phase for all four clusters. AMN-dominant states (Clusters 2 and 4) were preferentially expressed during planning, whereas DMN-dominant states (Clusters 1 and 3) were preferentially expressed during feedback. Phase profiles were therefore similar within each network family. By contrast, sensitivity to planning demand differed even when states shared the same dominant network. Among the DMN-dominant states, Cluster 3 occupancy decreased under higher demand whereas Cluster 1 was unchanged, and among the AMN-dominant states, Cluster 4 occupancy increased whereas Cluster 2 occupancy decreased. One possibility is that Cluster 4 is a more planning-sensitive AMN-dominant variant, but this remains a hypothesis. The presence of more than one AMN-dominant state with distinct demand sensitivity is broadly compatible with accounts in which cognitive control is implemented by multiple controllers^32^.

In CCTHot, among the DMN-dominant states, higher pre-decision occupancy of Cluster 3 was associated with continuing, whereas Cluster 1 showed no significant association. Among the AMN-dominant states, higher occupancy of Cluster 2 was associated with stopping, whereas Cluster 4 showed no significant association. As in WATT3, these results indicate that occupancy can relate to behavior in different ways even for states in the same DMN-or AMN-dominant family. Accumulating such separations across a broader range of cognitive tasks should help a more detailed understanding of these states.

### 3.5 Limitations

Several limitations define the scope of these conclusions. Our results are correlational and do not address any causal relationship between brain state and behavior. For example, it is unclear whether occupying a particular state changes performance or whether performance and state are jointly driven by a third factor. Our analyses assigned each time point to a single state (hard labeling). Soft-labeling approaches that allow multiple states to coexist at each time point remain an alternative for future work. We used a fixed 6-s shift to account for the hemodynamic delay, but individual or regional differences in the hemodynamic response function remain possible. Finer temporal examination of these dynamics will require methods such as EEG or MEG. Finally, the present task battery was weighted toward cognitive-control paradigms, so validation in more diverse settings, including video games and other naturalistic tasks, will be important.

### 3.6 Conclusions

In summary, the inside-out analysis revealed complementary levels of organization in shared brain state dynamics. Within individuals, the contrast between the common DMN- and AMN-dominant states showed relatively consistent relationships with response speed and variability but a task-condition-dependent relationship with accuracy. Across individuals, switch rate captured a cross-task individual component associated with the speed–accuracy trade-off, whereas occupancy primarily reflected task context. The DMN- and AMN-dominant variants observed in the planning and value-based decision paradigms further showed that states within the same dominant-network family can serve distinct behavioral functions. Together, these findings establish a framework in which momentary state identity and switching between states relate to trial-to-trial fluctuations in performance and to stable individual differences in response strategy, with task context shaping which states occur and what they mean for behavior. Integrating this inside-out characterization with psychological constructs established through outside-in approaches will provide a more comprehensive understanding of the neural basis of cognition.

## 4. Methods

### 4.1 Participants and tasks

We analyzed the publicly available OpenNeuro dataset (ds004636, version 1.0.4), which comprised *N* = 110 healthy adults (71 female, 64.5%; mean age 23.8 ± 5.6 years) who completed a battery of cognitive tasks during fMRI^5,33,34^ (see Supplementary Tables S1–S2 for per-task sample sizes). The task battery was designed to measure self-regulation across multiple domains^5,33^. The present analyses included 10 cognitive tasks spanning attention, response inhibition, cognitive control, planning, value-based decision making, and rest (Figure 1 A). Six tasks required speeded responses on each trial: the attention network test (ANT; reporting the direction of a central arrow flanked by congruent or incongruent arrows), cued task switching (twoByTwo; judging the magnitude or the color of a digit according to a cue), the Stroop task (naming the ink color of a color word), the dot pattern expectancy task (DPX; responding to cue–probe pairs that differ in expectancy), the simple stop-signal task (stopSignal; withholding a speeded response when a stop signal appears), and the motor-selective stop-signal task (motorSelectiveStop; withholding only one of two responses when the stop signal appears). Three tasks involved planning or value-based decisions: a towers planning task (WATT3; rearranging balls across three pegs to reach a target configuration), delay discounting (discountFix; choosing between a smaller immediate and a larger delayed reward), and the Columbia Card Task (CCTHot; turning over cards to accumulate points at the risk of drawing a loss card). The tenth condition was resting-state scans. Per-task sample sizes are reported in Supplementary Tables S1–S2. The shared dataset also includes a survey medley task (surveyMedley), which was excluded from the present analyses. Full task descriptions, including the conditions and trial types analyzed here, are provided in the Supplementary Information (Supplementary Methods, Cognitive tasks).

This study analyzed a publicly available, fully de-identified dataset, and no new data were collected. The original data collection was approved by the Stanford University Institutional Review Board (protocol no. 39322), and all participants provided written informed consent to participation and to the public release of their deidentified data^33^. The protocol for the present secondary analysis was approved by the institutional ethics committee of the Graduate School of Information Science and Technology, The University of Tokyo (approval no. 23-572). Participant sex was reported in the shared dataset as self-identified categories (71 female, 38 male, and 1 not reported); sex and gender were not analyzed as variables in the present study because our hypotheses concerned task-general state dynamics, and the sample was not designed to support sex- or gender-stratified inference.

fMRI data were acquired on a 3T GE Discovery MR750 scanner with a 32-channel head coil, using a single-echo multiband echo-planar imaging sequence (repetition time [TR] = 680 ms, echo time = 30 ms, flip angle = 53°, multiband factor = 8, 2.2 mm isotropic voxels, 64 slices, field of view = 220 mm); a high-resolution T1-weighted image (0.9 mm isotropic) was also acquired for registration.

Scans were excluded on a scan-by-scan basis for MRI quality (50 scans from 10 participants) or behavioral performance (19 scans from 16 participants) using the predefined criteria released with the shared dataset (Supplementary Table S3). After quality control and preprocessing, the number of participants entering brain state analyses ranged from 90 to 99 per cognitive task (Supplementary Table S1). Sample sizes for downstream analyses depended on their data-completeness requirements (Supplementary Table S2). Analyses of the within-individual consistency of state metrics included 66 participants with complete data for all nine non-rest tasks, whereas between-individual behavioral analyses included 91 participants who had state-dynamics metrics for at least four tasks and complete behavioral measures.

### 4.2 fMRI preprocessing and network-level time-series extraction

The fMRI data were preprocessed with fMRIPrep version 21.0.1^35^. Briefly, each T1-weighted anatomical image was corrected for intensity non-uniformity, skull-stripped, and spatially normalized to the MNI152NLin6Asym template. Each functional run underwent head-motion and susceptibility-distortion correction, was co-registered to the corresponding anatomical image, and was resampled to the same standard space at 2 mm isotropic resolution.

All subsequent signal processing was performed in Python using Nilearn. The preprocessed functional images were first spatially smoothed with a 6 mm full-width-at-half-maximum Gaussian kernel. Regional blood-oxygen-level-dependent (BOLD) time series were then extracted for the 400 cortical parcels of the Schaefer atlas^36^. During time-series extraction, nuisance regressors generated by fMRIPrep, including six head-motion parameters and the first six anatomical CompCor components^37^, were regressed out together with the global signal. The resulting parcel-wise time series were linearly detrended and standardized (z-scored) within each run.

Following the procedure used in our previous work^14^, we grouped the 400 parcels according to the seven canonical networks of the Yeo atlas^38^. Because the frontoparietal control network (FPCN) comprises two functionally distinct subsystems, one more strongly coupled with the default-mode network and the other with the dorsal-attention network^39^, we further divided its parcels into FPCN-A and FPCN-B according to their Yeo 17-network assignments^36^. Parcel-wise signals were then averaged within each of the resulting eight networks, yielding eight network-level time series per run (Figure 1 B). These time series served as the input to the brain state estimation described below.

### 4.3 Brain state definition and cross-task clustering

Following an inside-out strategy, we defined brain states separately for each of the 10 cognitive tasks as local minima in an energy landscape estimated from the fMRI signals, using the procedure established in previous studies^14,40,41^. Defining states separately in each task avoided imposing a common state structure a priori. For each task, we binarized the eight network-level time series at their temporal means and fitted a pairwise maximum entropy model (MEM) to the resulting binary activity patterns using variational expectation–maximization for the pairwise MEM (VEM-MEM;^42^). The model estimated a bias parameter for each network and pairwise interaction terms between networks. Under this model, the probability of a binary activity pattern ***s*** = (*s*_1_, … , *s*_8_) is *P*(***s***) ∝ exp[−*E*(***s***)], where the energy is

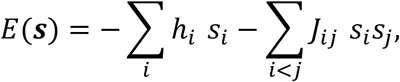

with ℎ*_i_* the bias of network *i* and *J_ij_* the pairwise interaction between networks *i* and *j* (see Supplementary Methods for parameter estimation). We then calculated the energy of each of the (2^8 = 256) possible binary activity patterns and identified local minima and their associated basins as brain states (Figure 1 C left).

To integrate these brain states across tasks while reducing dependence on the binary network representation, we derived a voxelwise gray-matter activation map for each state by averaging BOLD activity over the time points assigned to that state (Figure 1 C middle). We pooled the resulting maps across the 10 cognitive tasks and reduced their dimensionality using principal component analysis (PCA). A Gaussian mixture model (GMM) was then fitted to the scores on the first two principal components (PC1 and PC2), with the number of clusters (*k*) selected on the basis of the silhouette score across *k* = 2– 10 (Figure 1 C right). This procedure grouped the brain states obtained from each task into a common set of brain state clusters that occur across tasks. Each time point was thereby assigned to one of these clusters, yielding a cross-task brain state time series for each participant and task. As robustness checks, we repeated the clustering using (i) a higher-dimensional PCA subspace that retained 95% of the variance across the voxelwise maps and (ii) agglomerative hierarchical clustering of the PC1–PC2 scores (Supplementary Information).

To characterize the spatial pattern of each brain state cluster (Figure 2 D), we averaged the voxelwise activation maps of the brain states assigned to that cluster. Although brain states were estimated from cortical network time series only, these cluster-level maps covered the whole gray matter, and we summarized subcortical activity by averaging each map within the 16 regions of the Tian Subcortex S1 atlas^43^ (hippocampus, amygdala, anterior and posterior thalamus, nucleus accumbens, globus pallidus, putamen, and caudate, in each hemisphere). Values from the two hemispheres were averaged for display.

To describe state dynamics across the 10 cognitive tasks, we computed the occupancy of each brain state cluster and the transition probabilities between clusters for each participant and cognitive task, using all available scans for that task. Occupancy was defined as the proportion of time points assigned to each cluster, and transition probabilities were calculated between clusters at successive time points.

### 4.4 Within-individual associations between brain states and behavior

To investigate within-individual associations between brain states and performance, we focused on cognitive tasks characterized by repeated stimulus–response events (ANT, twoByTwo, Stroop, DPX, stopSignal, and motorSelectiveStop). For each task condition, we tested whether behavioral performance varied as a function of the occupied brain state cluster (Figure 1 E). Unless otherwise specified, each task event was associated with the brain state observed 6 s after event onset to account for the hemodynamic delay.

The behavioral outcomes were log-transformed reaction time (logRT), reaction-time variability quantified by the coefficient of variation (CV), and accuracy. LogRT and CV were analyzed using linear mixed-effects models, whereas trial-level accuracy was analyzed using a binomial generalized linear mixed-effects model. Each model included fixed effects of brain state, task condition, and their interaction, where applicable, together with a participant random intercept.

### 4.5 Between-individual differences in state dynamics and their behavioral correlates

#### 4.5.1 Cross-task consistency of individual state dynamics

State metrics can vary both between individuals and between tasks. To quantify how much of each source was present before relating state dynamics to individual behavior (Figure 1 F), we calculated the intraclass correlation coefficient ICC(3,1) in two complementary orientations. First, to quantify whether individual differences in each state metric were preserved across tasks (that is, whether participants were ordered similarly in every task), we treated participants as targets and tasks as raters. Second, to quantify whether differences among tasks were preserved across participants (that is, whether tasks were ordered similarly for every participant), we treated tasks as targets and participants as raters. As complementary measures of rank-order agreement, we also calculated pairwise Spearman correlations between tasks across participants and between participants across tasks.

#### 4.5.2 Associations with cross-task behavioral characteristics

To examine whether between-individual differences in state dynamics were associated with individual behavioral characteristics, we related the participant-level state-dynamics metrics described above to behavioral measures summarized across tasks (Figure 1 F). To extract the main factors underlying individual behavioral differences across tasks, we summarized accuracy and reaction-time measures with principal component analysis (PCA). Previous work has shown that such battery-wide measures can be decomposed into general performance and a speed–accuracy trade-off^5,19,20^. We applied PCA to 21 accuracy-related and 19 reaction-time-related measures defined at the task-and-condition level in ANT, twoByTwo, Stroop, DPX, stopSignal, and motorSelectiveStop.

Because not all participants completed all tasks, we computed task-averaged state-dynamics metrics as the mean of within-task *z*-scored values across available tasks, thereby reducing the influence of missing tasks. The primary sample included participants with state-dynamics metrics for at least four of the nine non-rest tasks. We calculated Spearman correlations between each participant’s behavioral component scores and each state-dynamics metric, with false discovery rate (FDR) correction across the two tests (PC1 and PC2) for each metric. The primary between-individual analyses focused on state-dynamics metrics that showed evidence of within-individual consistency across tasks in the participant-target ICC analysis. As a robustness analysis, we repeated the cross-task correlations in the 66 participants with complete data for all nine non-rest tasks.

### 4.6 Event-related state dynamics

Because cognitive control was a major focus of the task battery, several tasks included within-task conditions with well-defined differences in control demand. We used these contrasts to test whether brain state occupancy tracked moment-to-moment variation in cognitive control demands and the success or failure of response inhibition. Specifically, we compared incongruent with congruent trials in Stroop and ANT, switch with stay trials in twoByTwo, and failed with successful inhibition in stopSignal and motorSelectiveStop. For each contrast, we estimated occupancy time courses aligned to event onset. Unlike the trial-level state–behavior analyses, these analyses retained the full peri-event time course without applying a fixed 6-s shift.

### 4.7 Brain state clusters in value-based decision and planning paradigms

The preceding analyses focused on cognitive tasks characterized by repeated stimulus– response events. As a complementary analysis, we examined how brain state occupancy was related to decision and planning variables in discountFix, CCTHot, and WATT3. For each task and brain state cluster, we calculated mean occupancy within predefined task windows corresponding to trial phases, experimental conditions, or pre-decision periods, as detailed in the Supplementary Methods. We confirmed that every participant showed at least 1% occupancy of Clusters 3 and 4 in each of these paradigms (Supplementary Table S8).

In WATT3, a towers planning task, problems varied in planning demand, and each round comprised three phases: planning, action, and feedback ( ≈ 3 s, ≈ 3 s, and 1 s). We examined whether occupancy of each brain state cluster differed between lower-and higher-demand problems and across these phases. Mean occupancy was calculated for each round and phase and analyzed using two linear mixed-effects models, one with fixed effects of planning demand, cluster, and their interaction, and one with fixed effects of trial phase, cluster, and their interaction, together with a participant random intercept.

In CCTHot, participants turned over cards one by one from an array of multiple cards that included both wins and losses. Drawing a win added points, whereas drawing a loss ended the round, and on each decision they could stop or continue. To examine whether the choice to continue or stop differed with the immediately preceding brain state, we compared trials matched for expected value, which was determined by the remaining cards and the size of the loss. Because one-by-one choices were separated by only about 2 s and could not be resolved in the BOLD signal, we restricted this comparison to round-final decisions, matching each round-final stop decision to a continue decision from the same participant with a similar expected value. We used mixed-effects logistic regression with a participant random intercept to test whether mean occupancy over the seven TRs (≈ 4.8 s) preceding the decision was associated with the probability of continuing versus stopping.

In discountFix, participants chose on each trial between a smaller-sooner (SS) option that was always $20 today and a larger-later (LL) option whose amount and delay varied across trials (for example, $30 in 23 days). We focused on mid trials, in which the two options were selected at approximately equal rates and choice was therefore expected to be most difficult. We then asked whether the immediately preceding brain state was related to the upcoming choice. We used mixed-effects logistic regression with a participant random intercept to test whether mean occupancy over the seven TRs (≈ 4.8 s) preceding the decision was associated with the probability of an LL choice.

### 4.8 Use of AI-assisted tools

The authors used Cursor (https://cursor.com) to assist with language editing and implementation of the analysis code. The authors reviewed and revised the AI-assisted text as appropriate and reviewed and tested the AI-assisted code. All scientific decisions and interpretations were made by the authors, who take full responsibility for the accuracy and integrity of the manuscript, analysis code, and reported results.

## 5. Data availability

The fMRI and behavioral data analyzed in this study are openly available from OpenNeuro (ds004636, version 1.0.4) and are described in the corresponding data descriptor^33^.

## 6. Code availability

The analysis code is available at https://github.com/Ayumu722/multicog_brainstate (which will be made publicly available upon acceptance of the manuscript). We implemented this estimation in Python following the algorithm and the original reference implementation developed by Jeong, Kang, and colleagues^41,42^.

## 7. Acknowledgements

This work was supported by the Japan Agency for Medical Research and Development (AMED) under Grant Numbers JP25wm0625417 and JP25wm0625219. The funder had no role in study design, data collection and analysis, decision to publish, or preparation of the manuscript. We thank the investigators who collected and shared the OpenNeuro ds004636 dataset.

## 8. Author contributions

A.Y. conceived and designed the study, performed the analyses, and wrote the manuscript. M.S. contributed to the interpretation of the results and revised the manuscript. K.A. contributed to the interpretation of the results and revised the manuscript.

## References

1. Posner, M. I., Petersen, S. E., Fox, P. T. & Raichle, M. E. Localization of cognitive operations in the human brain. *Science (New York*, N.Y.) 240, 1627–1631 (1988).

2. Poldrack, R. A. Can cognitive processes be inferred from neuroimaging data? Trends in Cognitive Sciences 10, 59–63 (2006).

3. Nau, M., Schmid, A. C., Kaplan, S. M., Baker, C. I. & Kravitz, D. J. Centering cognitive neuroscience on task demands and generalization. Nature neuroscience 1–12 (2024) doi:10.1038/s41593-024-01711-6.

4. Poldrack, R. A. & Yarkoni, T. From brain maps to cognitive ontologies: Informatics and the search for mental structure. Annual review of psychology 67, 587–612 (2016).

5. Eisenberg, I. W. et al. Uncovering the structure of self-regulation through data-driven ontology discovery. Nature communications 10, 2319 (2019).

6. Nakuci, J. et al. Multiple brain activation patterns for the same perceptual decision-making task. Nature Communications 16, 1785 (2025).

7. Buzsáki, G. The Brain from Inside Out. (Oxford University Press, Cary, NC, 2019). doi:10.1093/oso/9780190905385.001.0001.

8. Greene, A. S., Horien, C., Barson, D., Scheinost, D. & Todd Constable, R. Why is everyone talking about brain state? Trends in neurosciences 0, (2023).

9. Vidaurre, D., Smith, S. M. & Woolrich, M. W. Brain network dynamics are hierarchically organized in time. Proceedings of the National Academy of Sciences of the United States of America 114, 12827–12832 (2017).

10. Wu, Y.-H., Podvalny, E., Levinson, M. & He, B. J. Network mechanisms of ongoing brain activity’s influence on conscious visual perception. Nature communications 15, 5720 (2024).

11. Song, H., Shim, W. M. & Rosenberg, M. D. Large-scale neural dynamics in a shared low-dimensionalstate space reflect cognitive and attentional dynamics. eLife 12, (2023).

12. Cai, W., Taghia, J. & Menon, V. A multi-demand operating system underlying diverse cognitive tasks. Nature communications 15, 2185 (2024).

13. Yun, X., Cai, W., Fun, J., Zhu, X. & Zhao, X. State-space modeling uncovers brain-behavior dynamics of inhibitory control. Cerebral Cortex (New York, N.Y.: 1991) 35, bhaf197 (2025).

14. Yamashita, A., Rothlein, D., Kucyi, A., Valera, E. M. & Esterman, M. Brain state-based detection of attentional fluctuations and their modulation. NeuroImage 236, 118072 (2021).

15. Yamashita, A. et al. Variable rather than extreme slow reaction times distinguish brain states during sustained attention. Scientific reports 11, 14883 (2021).

16. Pessoa, L., Medina, L. & Desfilis, E. Refocusing neuroscience: Moving away from mental categories and towards complex behaviours. Philosophical transactions of the Royal Society of London. Series B, Biological sciences 377, 20200534 (2022).

17. Shine, J. M. et al. Human cognition involves the dynamic integration of neural activity and neuromodulatory systems. Nature neuroscience 22, 289–296 (2019).

18. Dosenbach, N. U. F., Raichle, M. E. & Gordon, E. M. The brain’s action-mode network. Nature reviews. Neuroscience 26, 158–168 (2025).

19. Friedman, N. P. & Miyake, A. Unity and diversity of executive functions: Individual differences as a window on cognitive structure. Cortex; a Journal Devoted to the Study of the Nervous System and Behavior 86, 186–204 (2017).

20. Hedge, C., Powell, G., Bompas, A. & Sumner, P. Strategy and processing speed eclipse individual differences in control ability in conflict tasks. Journal of Experimental Psychology. Learning, Memory, and Cognition 48, 1448–1469 (2022).

21. Englert, R. et al. Functional connectivity-based attractor dynamics of the human brain in rest, task, and disease. eLife 13, (2026).

22. Gözükara, D., Oetringer, D., Ahmad, N. & Geerligs, L. Multi-scale anti-correlated neural states dominate naturalistic whole-brain activity. eLife https://doi.org/10.7554/elife.109116.1 (2026) doi:10.7554/elife.109116.1.

23. Lawrence, N. S., Ross, T. J., Hoffmann, R., Garavan, H. & Stein, E. A. Multiple neuronal networks mediate sustained attention. Journal of cognitive neuroscience 15, 1028–1038 (2003).

24. Raichle, M. E. The brain’s default mode network. Annual review of neuroscience 38, 433–447 (2015).

25. Kahneman, D. Thinking, Fast and Slow. (macmillan, 2011).

26. Gronchi, G. & Giovannelli, F. Dual process theory of thought and default mode network: A possible neural foundation of fast thinking. Frontiers in psychology 9, 1237 (2018).

27. Vatansever, D., Menon, D. K. & Stamatakis, E. A. Default mode contributions to automated information processing. Proceedings of the National Academy of Sciences of the United States of America 114, 12821–12826 (2017).

28. Schneider, W. & Shiffrin, R. M. Controlled and automatic human information processing: I. Detection, search, and attention. Psychological review 84, 1–66 (1977).

29. Shenhav, A. et al. Toward a rational and mechanistic account of mental effort. Annual review of neuroscience 40, 99–124 (2017).

30. Amer, T., Campbell, K. L. & Hasher, L. Cognitive control as a double-edged sword. Trends in cognitive sciences 20, 905–915 (2016).

31. Hearne, L., Cocchi, L., Zalesky, A. & Mattingley, J. B. Interactions between default mode and control networks as a function of increasing cognitive reasoning complexity. Human brain mapping 36, 2719–2731 (2015).

32. Dosenbach, N. U. F., Fair, D. a., Cohen, A. L., Schlaggar, B. L. & Petersen, S. E. A dual-networks architecture of top-down control. Trends in cognitive sciences 12, 99–105 (2008).

33. Bissett, P. G. et al. Cognitive tasks, anatomical MRI, and functional MRI data evaluating the construct of self-regulation. Scientific Data 11, 809 (2024).

34. Eisenberg, I. W. et al. Applying novel technologies and methods to inform the ontology of self-regulation. Behaviour research and therapy 101, 46–57 (2018).

35. Esteban, O. et al. fMRIPrep: A robust preprocessing pipeline for functional MRI. Nature methods 16, 111–116 (2019).

36. Schaefer, A. et al. Local-global parcellation of the human cerebral cortex from intrinsic functional connectivity MRI. Cerebral cortex (New York, N.Y.: 1991) 28, 3095–3114 (2018).

37. Behzadi, Y., Restom, K., Liau, J. & Liu, T. T. A component based noise correction method (CompCor) for BOLD and perfusion based fMRI. NeuroImage 37, 90–101 (2007).

38. Yeo, B. T. T. et al. The organization of the human cerebral cortex estimated by intrinsic functional connectivity. Journal of neurophysiology 106, 1125–1165 (2011).

39. Dixon, M. L. et al. Heterogeneity within the frontoparietal control network and its relationship to the default and dorsal attention networks. Proceedings of the National Academy of Sciences 115, E1598–E1607 (2018).

40. Ezaki, T., Watanabe, T., Ohzeki, M. & Masuda, N. Energy landscape analysis of neuroimaging data. Philosophical transactions. Series A, Mathematical, physical, and engineering sciences 375, (2017).

41. Kang, J., Jeong, S.-O., Pae, C. & Park, H.-J. Bayesian estimation of maximum entropy model for individualized energy landscape analysis of brain state dynamics. Human Brain Mapping 42, 3411–3428 (2021).

42. Jeong, S.-O. et al. Empirical bayes estimation of pairwise maximum entropy model for nonlinear brain state dynamics. NeuroImage 244, 118618 (2021).

43. Tian, Y., Margulies, D. S., Breakspear, M. & Zalesky, A. Topographic organization of the human subcortex unveiled with functional connectivity gradients. Nature neuroscience 23, 1421–1432 (2020).

