## Supplementary Information for "Task-general brain state dynamics relate to trial-to-trial performance and individual response strategy"

Ayumu Yamashita[1,2,3], Masataka Sawayama[4], Kaoru Amano[2]
[1]Graduate School of System Informatics, Kobe University, Kobe, Japan
[2]Department of Information Physics and Computing, Graduate School of Information Science and Technology, The University of Tokyo, Tokyo, Japan
[3]Brain Information Communication Research Laboratory Group, Advanced Telecommunications Research Institutes International, Kyoto, Japan
[4]Graduate School of Information Science and Technology, Hokkaido University, Sapporo, Japan

#### 1. Supplementary Figures

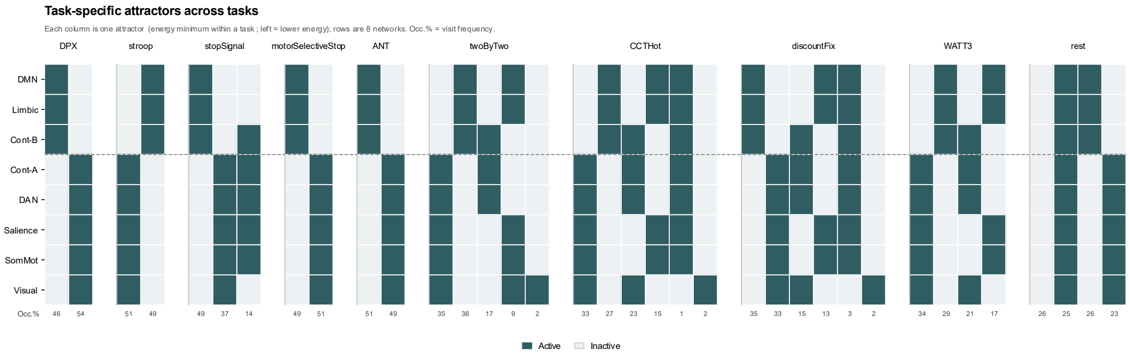


Figure S1: Energy landscape and brain states obtained from each task, identified as its local minima.

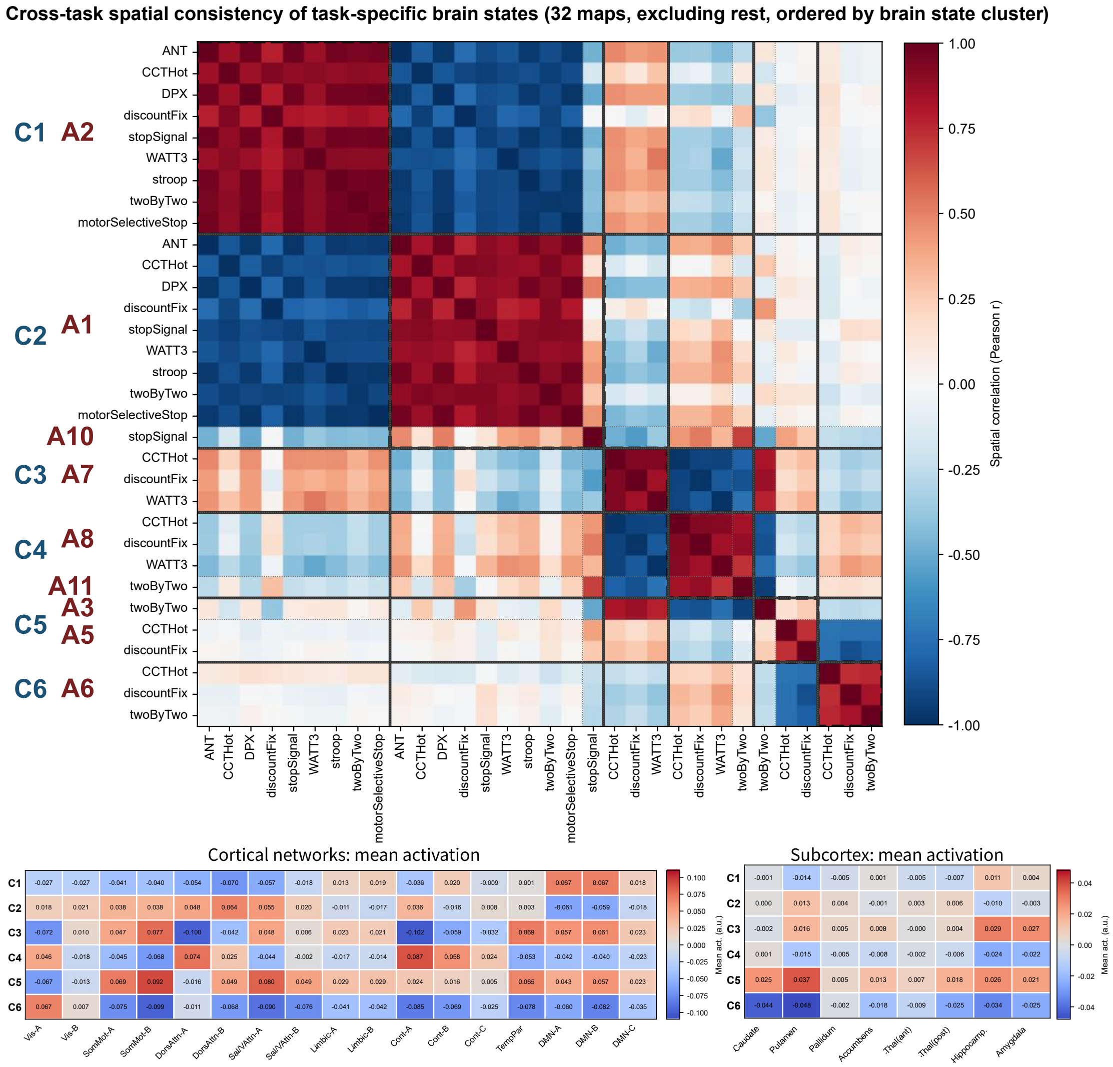


Figure S2: Spatial consistency of the seven brain state clusters across tasks.

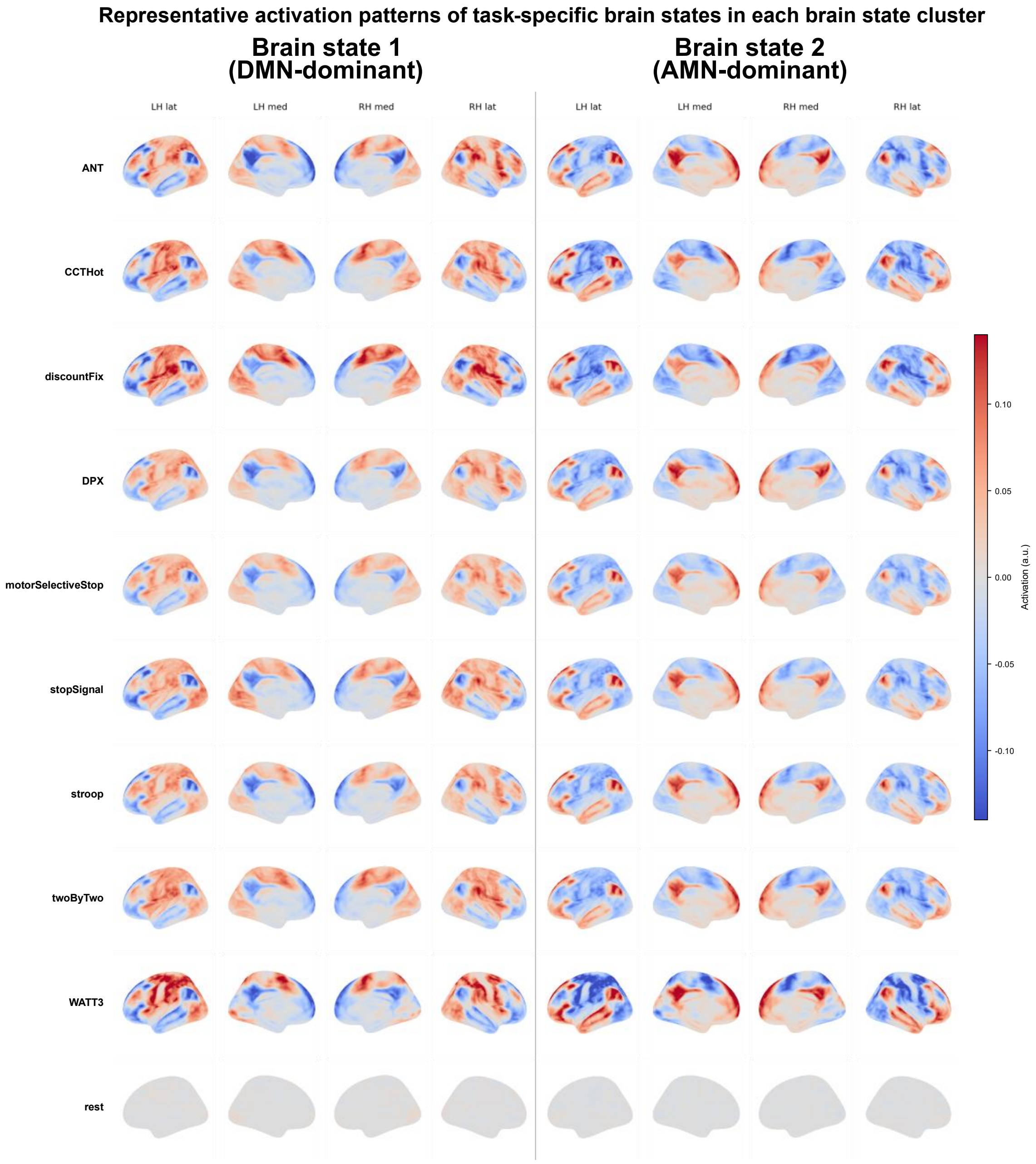


Figure S3: Representative activation patterns of the seven brain state clusters.

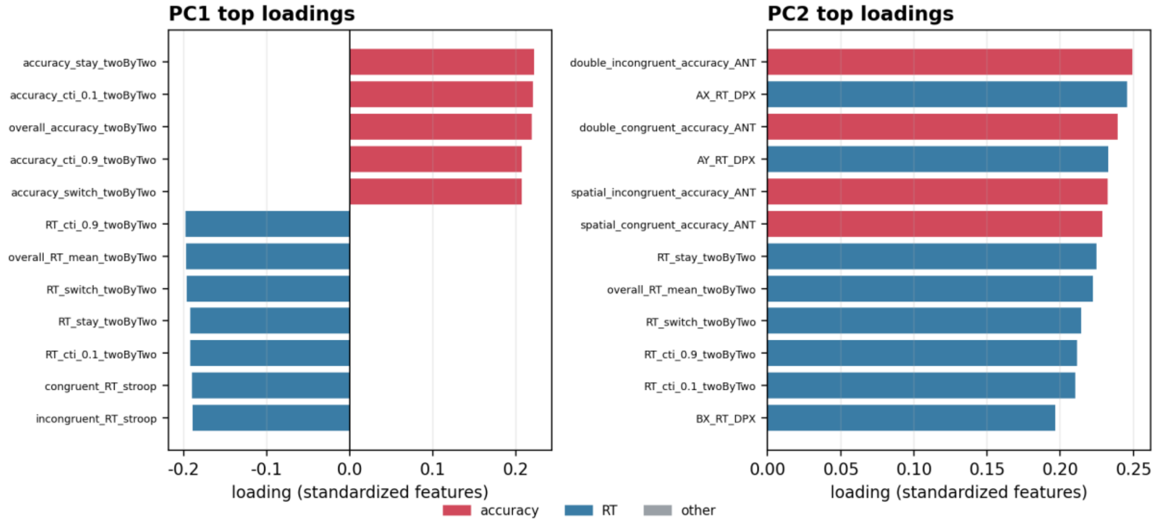


Figure S4: Interpretation of the behavioral principal components (PC1, general performance; PC2, speed–accuracy trade-off).

### 2. Supplementary Methods

#### 2.1 Dataset and participants

We analyzed an openly shared dataset acquired to study the construct of self-regulation, available through OpenNeuro (ds004636, version 1.0.4) and described in full in the accompanying data descriptor[1]. The summaries below paraphrase the aspects of the acquisition and tasks that are most relevant to the present study. Readers are referred to the original data descriptor for complete procedural detail. In brief, 110 healthy adults were enrolled in the shared cohort and completed up to two 90-minute scanning sessions on a 3 T scanner, during which they performed a battery of cognitive tasks together with resting-state and structural scans. The tasks had been selected from the larger behavioral battery of Eisenberg et al. using a data-driven procedure intended to capture the breadth of individual differences in self-regulation[2,3]. Functional images were acquired with a multiband echo-planar sequence (repetition time [TR] = 0.68 s, 2.2 mm isotropic voxels). Our analyses used the subsets of participants and tasks described in the main text (Table S1). Scans were excluded on a scan-by-scan basis according to the quality-control flags released with the shared dataset. MRI-quality exclusions were based on MRIQC failure (50 scans from 10 participants), and behavioral exclusions were based on a stop-success rate below 25% or above 75% in the stop-signal tasks, a non-critical omission rate above 35% in motorSelectiveStop, or a non-response rate above 50% in any task (19 scans from 16 participants in total).

#### 2.2 Cognitive tasks

Below we briefly summarize each in-scanner task. Each description follows the data descriptor[1], and the original paradigm references are given in the text.

**Attention network test (ANT).** Originally introduced by Fan et al.[4], this task probes alerting, orienting, and executive attention. On each trial a row of five arrows appears above or below fixation, and participants report the direction of the central arrow. The flanking arrows either point in the same direction as the central arrow (congruent) or in the opposite direction (incongruent). The target is preceded by a cue that is either non-informative (a double cue) or spatially predictive of the target location. The key manipulation for the present study is the congruency contrast, which indexes the resolution of response conflict. The task comprised 128 trials in two blocks.

**Cued task-switching (twoByTwo).** Following the cued task-switching paradigm[5,6], a central cue instructed participants on each trial to judge either the magnitude (greater or less than 5) or the color (orange or blue) of a subsequently presented colored digit. Because each task was indicated by two interchangeable cues, the design dissociates trials in which the cue repeats (cue stay), the cue changes but the task stays the same (cue switch), and the task itself changes (task switch). The task comprised 240 trials in three blocks.

**Columbia card task (hot version, CCTHot).** In this risk-taking paradigm[7], participants sequentially turned over cards to accumulate points, with each array specifying the number of loss cards and the magnitude of gains and losses. Turning over a gain card added points, whereas a loss card ended the round. Participants could stop (“cash out”) at any time. Because the round ended at a loss card, the task captures a balance between reward seeking and risk avoidance. Rounds were self-paced, with the run terminated after about 12 minutes.

**Dot-pattern expectancy task (DPX).** In this task[8,9], participants viewed cue–probe pairs formed from dot patterns and made a speeded response to each probe. One response was required when a designated target cue (A) was followed by a designated target probe (X), and an alternative response was required for all other pairings. The four conditions (AX, AY, BX, BY) differ in the expectancy set up by the cue, allowing proactive and reactive cognitive control to be distinguished. The task comprised 160 trials in four blocks, with AX trials most frequent.

**Delay-discounting task (discountFix).** On each trial[10], participants chose between a smaller immediate reward ($20 today) and a larger reward available after a delay, with the delayed amount and delay varying across trials. Choices index the degree to which future rewards are devalued by delay. The task comprised 120 trials in two blocks.

**Stop-signal task.** In this canonical measure of response inhibition[11], participants made speeded categorical responses to “go” shapes. On a subset of trials a stop signal appeared after a variable delay, instructing them to withhold the response. The stop-signal delay was adjusted trial-by-trial with a staircase so that stopping succeeded on roughly half of stop trials. The task comprised 125 trials, of which 40% were stop trials.

**Motor selective stop-signal task.** This variant[12] was identical to the stop-signal task except that the stop signal applied only to one ("critical") response. Participants were to withhold that response when the signal appeared but to continue responding when the signal followed the other ("non-critical") response. This isolates selective, motor-specific inhibition. The task comprised 250 trials (60% go, 20% critical stop, 20% non-critical stop).

**Stroop task.** In this classic interference task[13], a color word was printed in colored ink, and participants reported the ink color while ignoring the word. The word and ink color were either the same (congruent) or different (incongruent), and the incongruent-minus-congruent contrast indexes interference control. The task comprised 96 trials in two blocks.

**Towers task (Ward–Allport, WATT3).** In this planning task[14], participants rearranged balls across three pegs to reach a target configuration in as few moves as possible. Trials were designed to be solvable in three moves and varied in whether an intermediate move was required, manipulating planning demand. The task comprised up to 48 self-paced rounds.

**Survey medley.** Participants also answered 40 self-report items presented in the scanner, drawn from established self-regulation questionnaires. This non-speeded condition was not a focus of the present analyses.

#### 2.3 Energy-landscape analysis and clustering

##### 2.3.1 Binarization of network activity

For each scan, the time series of the eight networks were first converted into a sequence of binary activity patterns. Each network signal was standardized (z-scored) over time. At every time point a network was labeled as active (1) when its standardized signal exceeded its own temporal mean and as inactive (0) otherwise. With eight networks, this binarization maps the data onto one of $2^8 = 256$ possible activity patterns at each time frame. The empirical data are summarized by how frequently each network is active and how often pairs of networks are co-active.

##### 2.3.2 Pairwise maximum entropy model

We then described the distribution of binary patterns with a pairwise maximum entropy model (MEM). Among all probability distributions that reproduce the observed mean activity of each network and the observed co-activation of every pair of networks, the MEM is the one that is otherwise maximally unstructured. That is, the distribution adds no statistical dependency beyond these first- and second-order constraints[15]. Under this model the probability of an activity pattern $\boldsymbol{s} = (s_1, \ldots, s_8)$, with $s_i \in \{0,1\}$ (active = 1, inactive = 0), takes the Boltzmann form $P(\boldsymbol{s}) \propto \exp[-E(\boldsymbol{s})]$, where the energy is

$$E(\boldsymbol{s}) = -\sum_{i} h_i \; s_i - \sum_{i<j} J_{ij} \; s_i s_j.$$

Here the bias term $h_i$ reflects the baseline tendency of network $i$ to be active, and the interaction term $J_{ij}$ reflects the tendency of networks $i$ and $j$ to be co-active. Patterns with low energy occur frequently, whereas high-energy patterns are rare.

##### 2.3.3 Parameter estimation by variational expectation–maximization

Rather than fitting the bias and interaction parameters by conventional maximum likelihood, which can be unstable for short individual recordings, we estimated them with the variational expectation–maximization scheme for the pairwise MEM (VEM-MEM) introduced by[16]. In this empirical-Bayes formulation the parameters are treated as latent variables endowed with prior distributions. Their posterior estimates are obtained by alternating between an expectation step and a maximization step until convergence. The model is hierarchical, estimating both a task-level (population) set of parameters and subject-specific parameters. This regularization yields more reliable parameters than point estimation, particularly when the amount of data per task is limited[16,17].

##### 2.3.4 Energy landscape and brain state definition

For each task, we constructed a single energy landscape from the task-level (population) parameters, so that brain states are defined at the group level per task. The fitted energy function assigns an energy value to each of the 256 activity patterns, defining an energy landscape over the discrete pattern space. Two patterns were regarded as neighbors when they differed in the state of a single network. A pattern was identified as a local minimum (a brain state) when its energy was lower than that of all of its neighbors, so that no single network flip could further decrease the energy. These local minima correspond to stable, recurring activity configurations. The remaining patterns were assigned to the basin of the brain state reached by iteratively moving to the lowest-energy neighbor. Each participant's binarized time series was in turn assigned to these group-level basins[15,18].

##### 2.3.5 Cross-task integration and clustering

Brain states were obtained per task as the local minima of the binarized energy landscape. To reduce the influence of the binarization step on the across-task comparison, we did not cluster the binary patterns directly. Instead, for each brain state we computed the corresponding voxel-level activation pattern in gray matter, which preserves the continuous spatial detail of the state. This yielded 36 (task × brain state) maps across the 10 tasks.

To integrate these maps across tasks, we reduced their dimensionality with principal component analysis (PCA) and clustered them with a Gaussian mixture model (GMM) in the plane defined by the first two principal components (PC1 and PC2). We selected the number of clusters $k$ on the basis of the silhouette score across $k = 2$–$10$. As robustness checks, we also clustered the maps (i) in the 7-dimensional PCA subspace that retained 95% of the variance and (ii) with agglomerative hierarchical clustering of the PC1–PC2 scores.

##### 2.3.6 Clustering results and robustness

Applying this procedure yielded seven brain state clusters (Clusters 0–6). PC1 and PC2 explained 42% and 29% of the variance across maps, respectively (71% cumulatively), and $k = 7$ was selected by the silhouette score.

The grouping of the 36 (task × brain state) maps into seven brain state clusters was robust to the clustering algorithm. Replacing the GMM with agglomerative hierarchical clustering of the same PC1–PC2 representation recovered the same partition (adjusted Rand index = 1.000; 0 of 36 maps reassigned). Repeating the clustering in the 7-dimensional PCA subspace that retained 95% of the variance yielded a closely related partition (adjusted Rand index = 0.946 relative to the PC1–PC2 GMM). However, the silhouette-optimal solution had eight clusters because one stopSignal map formed a cluster of its own.

#### 2.4 Brain state clusters in value-based decision and planning paradigms

The main text reports complementary analyses of decision and planning variables (choice, expected value, planning difficulty, and problem-solving phase) in discountFix, CCTHot, and WATT3. Below we specify the task windows, outcomes, and statistical tests. State labels followed the cross-task clustering described above (Clusters 0–6). These analyses used Clusters 1–4 because Clusters 3 and 4 were engaged in addition to the two common states in these tasks. As in the trial-level state–behavior analyses, but not the event-related occupancy time courses, the state time series was shifted by 6 s relative to event onsets to account for the hemodynamic delay.

##### 2.4.1 Task windows and behavioral coding

**WATT3 (planning).** Trials varied in planning demand. Some configurations could be solved by moving balls directly to their goal positions, whereas others required an *intermediate* move (temporarily relocating a ball that blocked the goal), which increases planning difficulty[1]. Each round was also segmented using BIDS event onsets into three phases: *planning* (from round onset until the first move), *action* (from the first move until the correct solution), and *feedback* (the post-solution feedback screen). For each participant, round, and phase, we computed the mean occupancy of Clusters 1–4 within that phase window.

**CCTHot (value-based decision).** Card-turn (flip) events defined the sequential decisions within each round. Expected value (EV) for each round was computed from the displayed gain and loss contingencies. Successive card decisions were too closely spaced for the BOLD signal to resolve individual decisions (median inter-decision interval about 2 s), so analyses aligned to every card turn were not interpretable at the decision level. We therefore compared the round-final decision to stop (cash out) with a continue decision (draw) matched within participants for EV (1:1 matching of decisions whose expected values differed by at most 1.0; decisions at card position 3 or later). We averaged state occupancy over the seven TRs (4.76 s at TR = 0.68 s) preceding each decision.

**discountFix (value-based decision).** Each trial comprised a 4-s offer presentation followed by a response period during which participants chose the smaller-sooner ($20 today) or larger-later (LL) option. We focused on *mid* trials (offers calibrated to yield approximately balanced LL and SS choices) and averaged occupancy over the seven TRs (4.76 s at TR = 0.68 s) immediately preceding the choice response (*pre-response window*, main analysis). The predictor was the continuous Cluster 1 minus Cluster 2 occupancy difference in this window.

##### 2.4.2 Statistical analysis

In WATT3, occupancy was analyzed at the round × phase level with linear mixed-effects models including participant random intercepts. We tested planning demand, cluster, and their interaction in one model (occupancy ∼ demand × cluster) and trial phase, cluster, and their interaction in a second model (occupancy ∼ phase × cluster). Simple effects of demand and Wald tests of the omnibus phase effect are reported per cluster. In CCTHot, we modeled the probability of continuing versus stopping from pre-decision occupancy of each cluster using mixed-effects logistic regression. In discountFix, we modeled the probability of an LL choice on mid trials from the continuous Cluster 1 minus Cluster 2 occupancy difference in the pre-response window using mixed-effects logistic regression with a participant random intercept. As a sensitivity analysis, we also modeled this occupancy difference across windows of 1 to 10 TRs (Table S2).

Main-text sample sizes were $n = 96$ (WATT3), $n = 89$ (CCTHot; EV-matched pairs), and $n = 94$ (discountFix). Relative to the per-task brain state samples with behavior (Table S1), these value-based and planning analyses, as well as the event-related analyses (Figure 5; e.g., $n = 96$ Stroop, $n = 93$ ANT, $n = 92$ twoByTwo, $n = 93$ stopSignal, $n = 87$ motorSelectiveStop), additionally excluded the two participants (sub-s635 and sub-s650) flagged as behavioral outliers for task non-compliance. Each task's sample is therefore two smaller than the corresponding behavioral-analysis $N$ in Table S1.

#### 3. Supplementary Tables

Table S1: Per-task sample flow from acquisition through quality control and preprocessing to the brain state and behavioral analysis samples. The shared dataset comprised 110 participants, of whom 108 had preprocessed fMRI time series. Resting-state scans had no behavioral events, so no acquisition-with-behavior or behavioral-analysis N is given. The surveyMedley task in the shared dataset was not analyzed.

| Task name | BIDS task | Acquired (fMRI + behavior) | MRI QC excluded | Behavioral QC excluded | QC pass with fMRI | Preprocessing incomplete | Brain state analysis *N* | Behavioral analysis *N* |
|---|---|---|---|---|---|---|---|---|
| Attention Network Test | ANT | 103 | 7 | 1 | 96 | 0 | 96 | 95 |
| Cued task switching | twoByTwo | 106 | 9 | 2 | 96 | 1 | 95 | 94 |
| Stroop task | stroop | 103 | 3 | 0 | 101 | 2 | 99 | 98 |
| Dot pattern expectancy task | DPX | 103 | 3 | 5 | 96 | 1 | 95 | 94 |
| Simple stop-signal task | stopSignal | 105 | 7 | 3 | 97 | 0 | 97 | 95 |
| Motor-selective stop-signal task | motorSelectiveStop | 104 | 4 | 8 | 93 | 3 | 90 | 89 |
| Towers planning task | WATT3 | 106 | 8 | 0 | 99 | 0 | 99 | 98 |
| Delay discounting | discountFix | 101 | 3 | 0 | 100 | 2 | 98 | 96 |
| Columbia Card Task | CCTHot | 104 | 7 | 0 | 99 | 0 | 99 | 97 |
| Resting-state | rest | – | 2 | 0 | 108 | 2 | 106 | – |

Additional QC and reconciliation tables (task scan availability, task disposition, MRI QC metrics, and verification that shared $N = 110$ reconciles with QC flags and fit logs) are available in the analysis repository under `paper_outputs/tables/` (table01c–table01g).

Table S2: discountFix sensitivity analysis: association between the continuous Cluster 1 minus Cluster 2 occupancy difference before the response and the probability of choosing the larger-later option on mid trials (mixed-effects logistic regression with participant random intercepts; 3,588 trials, $n = 94$). Pre-response windows are shown in TRs (TR = 0.68 s; main analysis: 7 TRs = 4.76 s). No window shows a significant association.

| pre-response window (TR) | ≈ s | $\beta$ | SE | $z$ | $p$ | OR |
|---|---|---|---|---|---|---|
| 1 | 0.7 | 0.044 | 0.059 | 0.75 | 0.45 | 1.04 |
| 2 | 1.4 | −0.000 | 0.063 | −0.00 | 1.00 | 1.00 |
| 3 | 2.0 | −0.015 | 0.067 | −0.23 | 0.82 | 0.99 |
| 4 | 2.7 | −0.016 | 0.071 | −0.23 | 0.82 | 0.98 |
| 5 | 3.4 | −0.038 | 0.074 | −0.51 | 0.61 | 0.96 |
| 6 | 4.1 | −0.042 | 0.076 | −0.54 | 0.59 | 0.96 |
| 7 | 4.8 | −0.053 | 0.079 | −0.68 | 0.50 | 0.95 |
| 8 | 5.4 | −0.049 | 0.081 | −0.60 | 0.55 | 0.95 |
| 10 | 6.8 | −0.031 | 0.086 | −0.36 | 0.72 | 0.97 |

### 4. References


1. Bissett, P. G. *et al.* Cognitive tasks, anatomical MRI, and functional MRI data evaluating the construct of self-regulation. *Scientific Data* **11**, 809 (2024).

2. Eisenberg, I. W. *et al.* Applying novel technologies and methods to inform the ontology of self-regulation. *Behaviour research and therapy* **101**, 46–57 (2018).

3. Eisenberg, I. W. *et al.* Uncovering the structure of self-regulation through data-driven ontology discovery. *Nature communications* **10**, 2319 (2019).

4. Fan, J., McCandliss, B. D., Fossella, J., Flombaum, J. I. & Posner, M. I. The activation of attentional networks. *NeuroImage* **26**, 471–479 (2005).

5. Logan, G. D. & Bundesen, C. Clever homunculus: Is there an endogenous act of control in the explicit task-cuing procedure? *Journal of Experimental Psychology. Human Peception and Performance* **29**, 575–599 (2003).

6. Mayr, U. & Kliegl, R. Differential effects of cue changes and task changes on task-set selection costs. *Journal of Experimental Psychology. Learning, Memory, and Cognition* **29**, 362–372 (2003).

7. Figner, B., Mackinlay, R. J., Wilkening, F. & Weber, E. U. Affective and deliberative processes in risky choice: Age differences in risk taking in the columbia card task. *Journal of Experimental Psychology. Learning, Memory, and Cognition* **35**, 709–730 (2009).

8. MacDonald, A. W. *et al.* A convergent-divergent approach to context processing, general intellectual functioning, and the genetic liability to schizophrenia. *Neuropsychology* **19**, 814–821 (2005).

9. Otto, A. R., Skatova, A., Madlon-Kay, S. & Daw, N. D. Cognitive control predicts use of model-based reinforcement learning. *Journal of Cognitive Neuroscience* **27**, 319–333 (2015).

10. Kable, J. W. *et al.* No effect of commercial cognitive training on brain activity, choice behavior, or cognitive performance. *The Journal of Neuroscience: The Official Journal of the Society for Neuroscience* **37**, 7390–7402 (2017).

11. Logan, G. D. & Cowan, W. B. On the ability to inhibit thought and action: A theory of an act of control. *Psychological Review* **91**, 295–327 (1984).

12. Aron, A. R., Behrens, T. E., Smith, S., Frank, M. J. & Poldrack, R. A. Triangulating a cognitive control network using diffusion-weighted magnetic resonance imaging (MRI) and functional MRI. *The Journal of Neuroscience: The Official Journal of the Society for Neuroscience* **27**, 3743–3752 (2007).

13. Stroop, J. R. Studies of interference in serial verbal reactions. *Journal of Experimental Psychology* **18**, 643–662 (1935).

14. Kaller, C. P., Rahm, B., Spreer, J., Weiller, C. & Unterrainer, J. M. Dissociable contributions of left and right dorsolateral prefrontal cortex in planning. *Cerebral Cortex (New York, N.Y.: 1991)* **21**, 307–317 (2011).

15. Ezaki, T., Watanabe, T., Ohzeki, M. & Masuda, N. Energy landscape analysis of neuroimaging data. *Philosophical transactions. Series A, Mathematical, physical, and engineering sciences* **375**, (2017).

16. Jeong, S.-O. *et al.* Empirical bayes estimation of pairwise maximum entropy model for nonlinear brain state dynamics. *NeuroImage* **244**, 118618 (2021).

17. Kang, J., Jeong, S.-O., Pae, C. & Park, H.-J. Bayesian estimation of maximum entropy model for individualized energy landscape analysis of brain state dynamics. *Human Brain Mapping* **42**, 3411–3428 (2021).

18. Yamashita, A., Rothlein, D., Kucyi, A., Valera, E. M. & Esterman, M. Brain state-based detection of attentional fluctuations and their modulation. *NeuroImage* **236**, 118072 (2021).
